# Four chromosomally clustered class-C β-lactamases (blaAmpH1/2/3/4) control the cephalosporin hydrolysis in *Mycobacterium tuberculosis* and similar genetic loci appeared as pseudogenes in *Mycobacterium leprae*

**DOI:** 10.1101/2023.09.04.556143

**Authors:** Asit Kumar Chakraborty

**Affiliations:** Department of Biotechnology and Biochemistry, Oriental Institute of Science and Technology-West Bengal, Vidyasagar University, India

**Keywords:** PBP4, blaAmpH, blaC, TB, cephalosporin, Leprosy, ISKpn transposase, BLAST- 2, AmpH primers

## Abstract

*Mycobacterium tuberculosis* (*Mt*) chromosomal ten PBPs and one class-A β-lactamase (blaC) were implicated in multi-drug resistance against penicillin, cephalosporin and carbapenem drugs. The *E. coli* Class-A PBP1A and PBP1B proteins referred as PonA1 and PonA2 in *Mt* with 34% homologies whereas Class-B PBP2 referred as PbpA and PbpB sub-class proteins with 27% homologies. The class-C PBPs were, PBP4, MecA_N, DacB1, DacB2 and AmpH1- AmpH4. During database search, we found that many *Mt* Pbp4 had no homology to *E. coli* PBP4 and such enzyme were found to be AmpH class with AmpC, ACC, DHA and GES β- lactamases similarities although one of such enzymes suggested as AmpH but the other referred as DacB or PBP5 in the literature. We investigated the *Mt* whole genomes (accession nos. AL123456, CP001642, CP054013, CP001641, CP025597) to get authentic PBP4 in *Mt* strain FDAARGOS_757 genome (protein id. AUP69687, 34% homology to *E. coli* PBP4) which was designated as conserved protein in chromosomes of *Mt* strains H37Rv, H37Rv-1, GG-36-11 and CCDC5180 making confusion in data analysis. Similarly, we BLASTP homology searched with plasmid-mediated 24 β-lactamases suggested that four *bla*AmpH genes, (*AmpH1, AmpH2, AmpH3* and *AmpH4*) predominantly control the degradation of cephalosporins in *M. tuberculosis* but such genes were found as pseudogenes in *M. leprae*. The *Mt* AmpH1/2/3/4 enzymes had better homologies with *V. parahaemolyticus* and *Yersinia pekkanenii* AmpH enzyme as well as with *E. coli* AmpH enzyme. The DacB1 (PBP6) and DacB2 (PBP7) enzymes had blaTEM similarity but no AmpH similarity indicating blaC and DacB1/B2 controlled the Penicillin hydrolysis in *Mt*. The blaC enzyme had 30% homology to *S. aureus* blaZ β-lactamase but such enzyme was also missing in *M. leprae*. Homology search suggested that carbapenem hydrolysis by blaOXA-23/51-like PBPA/B enzymes and blaOXA-58 related MecA_N domain β-lactamase which has 21% similarity to *S. aureus* mecA enzyme. The PonA1/A2 had no homology to 24 classes of β-lactamases but still were popular enzymes implicated for better penicillin hydrolysis. We designed primers for *Mt* PBPs to check transcription of individual genes by RT-PCR as well as to check chromosomal locus by BLASTN search after WGS. The *E. coli* genome had no similarities to *bla*AmpH1/2/3/4 genes but a 17nt (5’- CCTTGGTGCCGTCGACC-3’) sequence found in pKEC-a3c plasmid of *C. fruendii* with homology to *bla*AmpH4 gene (nt. 1236-1252) and shared a homology with IS1182-like ISKpn6 transposase of plasmids. The oligonucleotides selected many *Mt* chromosomes and located conserved blaAmpH4 protein in all cases. However, D435E and F495V mutations in the *Mt* strain 5521 blaAmpH4 were evident and frameshift deletions located in *Mt* stain FDAARGOS_756 *bla*AmpH4 gene with no protein was made. Thus, mutations, deletions and rearrangements mediated by IS-elements were the driving force to make new *AmpH* genes in *M. tuberculosis*.

## Introduction

The estimated 1·9 million deaths from tuberculosis in 2020, 500 000 more than in the previous year [1]. The COVID-19 pandemic between 2020-2022 highly impacted TB treatment similar to HIV pandemic earlier [2]. In 2023, 4.5 million TB infections is a concern for India (https://www.who.int/teams/global-tuberculosis-programme/data2023). The proper diagnosis is important for any disease to know when the disease has started and how it is progressing in the population. TB bacillus from patients hardly grow in medium-agar plate and thus drug sensitivity test is not possible to see which antibiotic is suitable to clear pathogen from host body. Eventually, all TB patients have to take 4-6 different drugs every day for at least 2 months and then 2-3 drugs for more 4 months [3–5]. In truth, within two weeks greater than 99% bacilli were dead but few slow growing and bone or deep tissue penetrated TB bacilli still remain and relapse will be the case if you do not continue 6 months multi-drug regime. Whatever the case, PENEM drugs were not popular for the treatment of TB although some popular high growing *Mycobacterium* strains were very sensitive to imipenem and meropenem [6–10]. The early TB cases (1940-1960) very easily cured by rifampicin and streptomycin drugs but *rpoB, rrs, rpsL* genes mutation caused drug resistance. The rifampicin inhibits RNA Polymerase enzyme by binding to it and RNA synthesis is blocked while β-subunit (rpoB gene) mutation gives resistance [11,12]. Some acetyl and phospho- transferases also caused streptomycin and aminoglycoside resistant due to acetylation and phosphorylation of the antibiotics that were protein synthesizing inhibitors [13,14]. The fluroquinolone drugs like ciprofloxacin, norfloxacin and moxifloxacin were also helpful to clear TB infection but DNA gyrase gene (gyrA/gyrB subunits) mutations caused drug resistance in many cases [15]. Macrolide antibiotics erythromycin, kanamycin and higher derivative amikacin were still used for the TB treatment but mmpL1-mmpL12 drug efflux genes and erythromycin esterase were implicated as their resistance [16–20]. Thus, all conventional drugs may not be helpful today and TB specific drugs like isoniazid, dapsone, ethambutol and linezolid were prescribed although some resistance to those very important drugs were already noticed through WGS sequencing [21–27]. In truth, no plasmid has been discovered in *M. tuberculosis* (*Mt*). Since 2015 few plasmids have been sequenced from other high replicating *Mycobacterium* strains but hardly found any *mdr* gene [28,29]. But multiple drugs intake is toxic and may be the cause of multi-drugs resistance TB but careful monitoring plasmid isolation must be followed. As for example, one multi-drug resistant *E. coli* strain KT-1 isolated from Ganga River at Kolkata and appeared no single plasmid in 1% agarose gel plus EtBr but finally it was found by pulse-gel-electrophoresis that many multiple sizes plasmids were giving smear and it was impossible to characterize all plasmids in such clone. Alternately, MDR bacteria isolated from chicken meat or milk (1.5ml miniprep) did not showed any plasmid from EtBr stained Agarose gel and some minor DNA in the lanes appeared as contaminated chromosomal DNA. But in reality, when four minipreps (6ml overnight culture) combined and loaded into gel, very distinctive medium size 27-30kb as well as low molecular weight 3kb and very high molecular weight >100-200kb plasmids bands appeared [30, 31]. Transformation experiment with *E. coli* DH5α calcium^+2^-cells with the excised plasmid gave distinctive antibiotic resistant colony, demonstrating the *amp* gene (ampicillin resistant) located in one plasmid and *cat* gene (chloramphenicol resistant) located in another larger plasmid [32]. In an elegant experiment with Indian TB patients and their family, it was concluded that nutritional food supplement caused rapid cure of TB patients as well as reduced the infections to other family members. Thus, we undertake to find the cause of penicillin drug resistance in TB and also want to find the cause of dissimilarity between *Mt* PBP4 verses *E. coli* PBP4 as we have seen in our study. We collected the all PBPs or β- lactamases from *Mt* genome by naked eye (5 hrs/genome; 4.4 million bases) and then compared with *E. coli* PBP1A, PBP1B, PBP2, PBP3, PBP4, PBP5, PBP6, PBP7 and AmpH as well and class-A, class-C and class-D β-lactamases like blaTEM, blaSHV, blaCTX-M, blaOXA-23, blaOXA-58, blaAmpC, blaKPC and blaNDM-1 to classify *Mt* PBPs and β-lactamases [33–36]. First antibiotic discovered in 1926 is penicillin from fungus *Penicillium notatam* through the process of fermentation [37]. Sadly, a penicillinase reported as early as 1940 and the said gene (amp) was detected in plasmid pBR322 during 1960s [38]. Further appearance of blaTEM, blaSHV, blaOXA penicillinases, scientist developed cephalosporin antibiotics with six atoms thiazole ring [33]. Sadly, *blaCTX-M-1/3/9* genes developed in plasmids of many enterobacteria to degrade cefotaxime and related cephalosporins (1970s) and scientist prepared synthetic carbapenem drugs (1980s) that were very popular now for extensive-drug-resistant pathogens [36]. Soltys discovered in 1952 that *M. tuberculosis* avian type grown in Dubos medium was more sensitive to penicillin than bovine and human types whereas *M. phlei* and BCG strains were the least sensitive [6]. Kasik and Peacham in 1968 reported that *M. smegmatis* (N.C.T.C. 8158), *M. fortuitum* and *M. phlei* (MPI) produced a constitutive cell-bound β-lactamase that had penicillinase and cephalosporinase activities with different specificities to penicillin-G, ampicillin and early derivative of cephalosporins [7]. Although twenty blaTEM, blaKPC, blaNDM, blaOXA, blaCTX-M like penicillinase isomers were reported in different enterobacteria, only a chromosomal class-A penicillinase (*blaC*) with homology to *blaTOHO* and *blaTEM* genes was reported in MDR *Mt* [1]. However, few penicillin binding proteins (PBPs) were reported in different *Mycobacterium* chromosome including *Mt* [39–43]. Such PBPs have 25-30% homologies with other bacterial penicillin-binding proteins like *E. coli*. *E. coli* possesses three class-A PBPs (PBP1a=OSL52747, PBP1b=AJF05047 and PBP1c=AKM36075) and two class-B PBPs (PBP2/penicillin resistant protein=AAB40835 and PBP3/peptidoglycan-binding protein=CAA38861) and five Class-C *E. coli* PBPs (PBP4=CAA42070, PBP5=EDV83509, PBP6=EGI51157, PBP7=OAF94491 and AmpH=ANK05711) [44–46]. *M. tuberculosis* produces two class A PBPs (ponA1 and ponA2), two class B PBPs (PbpA and PbpB) and a lipoprotein sharing some motifs with the class B PBPs [41]. Six classes C type PBP contained in *Mt*: one Pbp4, one Pbp5(dacB) which wrongly described one AmpH class in the database, one Pbp6 (DacB1), one Pbp7 (DacB2) and a putative type-AmpH which we showed here to be four sub-classes. Our *Mt* database search pinpointed the misuse nomenclature of *Mt* PBPs throughout the PubMed and NCBI Database. We have proved that Rv3627c, Rv1730c, Rv1367 and Rv1497 enzymes belong to AmpH class (Rv1730c only described as AmpH and Rv3627c as DacB or Pbp5 in database; AN:AL123456). The AmpH is a bifunctional DD-endopeptidase and DD-carboxypeptidase.

which cleaves the cross-linked dimers tetrapentapeptide with efficiencies K2/k= 1,200 M^-1^S^-^ and removes the terminal D-alanine from muropeptides with a C-terminal D-Ala-D-Ala dipeptide. However, a weak β-lactamase activity reported (1/1,000 the rate obtained for AmpC) for nitrocefin as substrate and the enzyme could be inhibited by 40 μM cefmetazole [47]. It was reported that *E. coli* AmpH binds to penicillin-G, cefoxitin, cephalosporin-C, cefmenoxime, cefotaxime but poorly with ampicillin, amoxicillin and carbenicillin [48]. The DacB1 (Pbp6) and DacB2 (Pbp7) related to class-A blaTEM group β-lactamases. The Rv2864c is related to blaOXA-23 and Rv2864c related to blaOXA-58 likely better perform carbapenem hydrolysis. No PBP related to *E. coli* Pbp3 was found in *Mt* chromosome and Pbp4 wrongly described and authentic *Mt* PBP4 enzyme were found in the database. We pinpointed here that *M. tuberculosis AmpH* gene was not one in the chromosome but blaAmpH1, blaAmpH2, blaAmpH3 and blaAmpH4 four such class-C β-lactamases would be regulated the degradation of cephalosporins. We also found 17nt sequence from AmpH4 gene had homology to ISKpn6 transposase gene of plasmids indicating new gene creation was mediated by DNA rearrangement involving IS-elements. We also made primers for all *Mt* PBPs to check their presence and location in the chromosome after WGS. The PubMed search with “AmpH + Mycobacterium” did not produce any result indicating role of *AmpH* gene in *Mt* wa*s* never investigated carefully.

## Methods

The NCBI GenBank database (www.ncbi.nlm.nih.gov/nucleotide/protein) used to get proteins and nucleotides for PBPs and β-lactamases. The BLASTN used to search nucleotide homology and BLASTP used to search protein homology. Multiple alignment of proteins was done by MultAlign software and multi-alignment was done by CLASTAL-Omega software [49–51]. We found some extended PBPs at the NH2-terminus and likely due to usage of upstream ATG codon instead UTU and UUT initiation codons found in many *Mt* genes. We found authentic *Mt* PBP4 in strain FDAARGOS_757 (AN: CP054013/ protein id AUP68265). But our attempts to find *Mt* PBP3 was failed as BLAST search limited to 5000 sequences only instead search for 20000 sequences were possible few years ago. We used “AmpH + Mycobacterium” to search PubMed but no result indicating role of AmpH β-lactamase in *Mt* never investigated carefully. We searched 4.4 million bases of *Mycobacterium* chromosome by naked eye in the computer GenBank database and it took 5 hours. We did BLAST-2 search with DNA to find homology and then homology sequence searched by BLAST-X to get important protein homologues if present in the *Mycobacterium* chromosome. Once we got the protein isomer, we back compared their penetration in the database by BLAST-P search with increasing sequence parameter to 5000 instead 50. *Vibrio* species had two chromosomes and we had analysed both separately. The *E. coli* PBPs and β-lactamase proteins used as standard but sometime *K. pneumoniae, S. marcessens, S. aureus, M. leprae, M. intracellulare* genomes and proteins were compared.

## Result

We used *E. coli* PBPs protein ids for BLASTP similarity search with 25-30% similarity (>40% cover) with *Mt* PBPs (see, supplementary files S1-S9 for more analysis). The PonA1 (823aa), PonA2 (810aa), PBPA (491aa), PBPB (605aa), MecA_N (582aa), blaC (307aa), PBP4 (473aa), DacB1 (386aa), dacB2 (291aa), AmpH1 (532aa), AmpH2 (377aa), AmpH3 (428aa) and AmpH4 (517aa) were the PBPs and β-lactamases reported in the database (see, S2). We found GTG initiation codon for *PonA1*, *PBPB* and *PBP4* genes whereas TTG initiation codon for *AmpH1* and *AmpH4* genes (see, S1). The PBP1A and PBP1B were the major transpeptidases- transglycosylases similar to *M. tuberculosis* PonA1 and PonA2 penicillin-binding-proteins. The PBP2 was assigned as PBPA and PBPB in *M. tuberculosis* genome. The sequence of PBP3 of *E. coli* did not match to *Mt* chromosome but some author put PBPB as PBP3. The DacA, DacB and DacC of *E. coli* referred as DacB (PBP5), DacB1 (PBP6) and DacB2 (PBP7) in *M. tuberculosis*. However, we found a MecA-N enzyme was implicated as PBP5 and sometime an AmpH homologue referred as PBP5 in *Mt*. The table-1 showed the classification and localization of different PBPs and β-lactamases whereas figure-1 showed the circular chromosomal localization of those proteins. We first got those enzymes from genomes of *M. tuberculosis* strain CCDC5180 (AN:CP001642) and *M. tuberculosis* strain H37Rv (AN: AL123456) (table-1). Blast search however, could not find an *E. coli* PBP3 homologue. We did not find any homology between *E. coli* PBP4 and the published *M. tuberculosis* PBP4. Recently, Kumar et al demonstrated that ponA1 and ponA2 could hydrolyse carbapenem drugs much better than blaC β-lactamase which implicated as sole β-lactamase in *Mt*. Thus, we understood that there was a problem in database nomenclature of *Mt* PBPs. It appeared that all available sequences of *E. coli* β-lactamases never compared with PBPs of *M. tuberculosis* genome. The BLASTP homology search could not find any similarity to bacterial MBL β- lactamases like blaOXA-23, blaOXA-48, blaOXA-51, blaOXA-58, blaKPC-1, blaNDM-1, blaIMP-1, blaGIM-1, blaVIM-1, and blaGES with PBP3 and PBP4 of *E. coli.* As expected some homology to blaDHA or blaACC with AmpC β-lactamase (protein id. CCP44496) and blaTEM with blaC β-lactamase (protein id. CCP44847) were found. Neither, we found any similarity of *Mt* MBL-like protein (protein id. CCP46589) with any bacterial β-lactamases and likely it was an Alkyl-Sulfatase rather than metallo-β-lactamase as disclosed in the database. Similarly, we BLASTP homology searched *Mt* PBP4 (protein id. CCP44126) with many β- lactamases found 25% (60% cover) similarity with class-C β-lactamases (blaAmpC, blaDHA, blaAAC, blaCMY) suggesting such class-C PBP had some capacity to hydrolyse penicillin drugs. However, database described many PBP4 in *Mt* genome but it was not true as such PBP proteins designated as DacA (class-B-type) whereas no similarity to *E. coli* PBP5, PBP6, PBP7 and related AmpH (D,D-carboxypeptidase and D,D-endopeptidase) (table-2). Both *Mt* PBP4 and blaDHA-like proteins have class C β-lactamase similarities but only has 29% (60% cover) similarity between themselves suggesting other *Mt* PBP4 may be exist and we have problem in nomenclature. The *Mt* PbpA and PbpB have no similarity to PBP4 (protein id. CCP44496) and no similarity found with class-A (blaTEM and blaCTX-M) and class-C (blaAmpC and blaDHA) β-lactamases. Still, we found a little similarity (27%/11 % cover) with blaOXA-58, blaOXA-177 and blaOXA-555 but not with blaOXA-1, blaOXA-2, blaOXA-23, blaOXA-48, blaOXA-226 and blaOXA-454 and blaOXA-565 suggesting class-B PBPs may have little MBL activities. Whereas, *Mt* PBPB (protein id. CCP45666) had similarity to blaOXA-23 and blaOXA-58 (26%/14% cover; protein id. AET95879 and 40%/9% cover, protein id. AGC92783) and might show MBL activities hydrolysing carbapenems. Further, *Mt* Pbp6 and Pbp7 have 27% (37% cover) similarity to Amp (pBR322 plasmid-derived first *mdr* gene discovered giving resistant to penicillin-G) and blaTEM-1 proteins (protein ids. AAB59737 and AKA86566) but not to class-C blaAmpC or class-D blaNDM-1 or blaOXA-48 suggesting that PBP6/7 will hydrolyse ampicillin but not carbapenem drugs. The PbpB has no similarity to Amp (Penicillinase) or blaTEM-1 Class-A and Class-C blaAmpC β-lactamases reflecting PBP3 and PBP5/6/7 were different proteins. Interestingly, PBP6 and PBP7 have 47.7% (63% cover) similarity between them and designated as DacB1 and DacB2 (D,D-carboxy peptidases) whereas PBP5 as DacB and PBP4 as DacA class-C PBPs. The *Mt* DacB PBP5 (532aa) had also similarity to blaAmpC (26.7%/46% cover) and blaKPC-1 (26%/18% cover) but no similarity to blaNDM-1, blaOXA-1, blaOXA-2, blaOXA-23, BlaOXA-48, blaOXA-51 and blaOXA-58 and thus will hydrolyse cephalosporins. The PBP3 of *E. coli* (protein id. CAA38861, AN: X55034, nt. 7943-9709) had no similarity to PBP4 and surprisingly did not detected any sPBP3 gene sequence in *Mt* genomes (AN: CP044345, CO040688, CP089775, CP040691, CP025597, CP054013, CP110619 and CP089927). The disclosed PBP4 gene in the database actually blaAmpH genes and truly PBP4 also found in the database with difficult. We found protein ids AIH68780, CNW39949, WP_003899610, AEJ48551, WP_003902671, AGL29101, NKE28321, CEZ25322 and EGB26878, as true PBP4 of *M. tuberculosis*. Such PBP4 also found in *M. bovis* (protein id. AHM09405), *M. avium* (protein id. WP_003873261), *M. intracellulare* (protein id. AFC46805), *M. colombiense* (protein id. OBJ06628), *M. europaeum* (protein id. WP_085241428), *M. timonense* (protein id. GFG99282), AFJ33415, *M. gordonae* (protein id. OBK44162), and *M. liflandii* (protein id. VLL08577). Thus, after careful homology analysis, we found four *bla*AmpH genes regulating the cephalosporin hydrolysis in *M. tuberculosis* (see below). Interestingly, *AmpH1, AmpH2, AmpH3* and *AmpH4* full length genes had no similarities to *E. coli* (AN:LN832404), *S. aureus* (AN:BX571856), *S. pneumoniae* (AN:NZ_CP107038) and *V. cholerae* (Ch-1, AN:NZ_CP028892; Ch-2, AN:NZ_CP012998) chromosomes although protein-protein homology was detected. Thus, we assume that *E. coli* has only one *AmpH* gene that has close similarity to other bacterial species (see later).

**Fig. 1.**
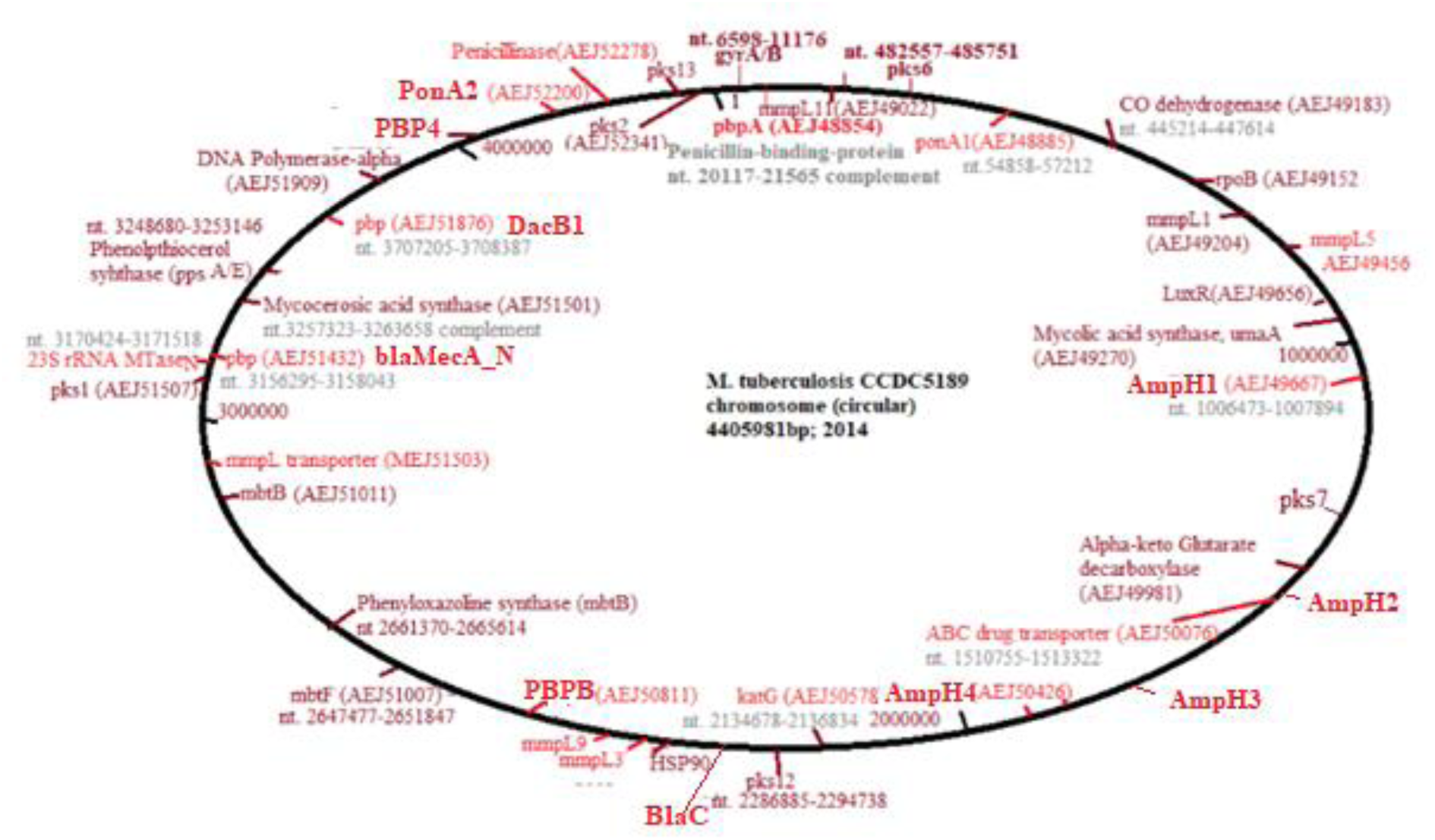
Localization of PBPs, blaC class-A β-lactamase and AmpH1/2/3/4 class-C β-lactamases including other important genes on the *M. tuberculosis* chromosome.

**Table-1:** Localization of penicillin-binding-proteins and Beta-lactamases in M. tuberculosis CCDC5180 genome (acc. no. CP001642) and compared with M. tuberculosis H37Rv genome (acc. no. AL123456) genome.

| **Table-1: Localization of penicillin-binding-proteins and Beta-lactamases in *M. tuberculosis* CDC5180 genome (acc. no. CP001642) and compared with *M. tuberculosis* H37Rv genome (acc. no. AL123456) genome** |  |  |  |
| --- | --- | --- | --- |
| Name of PBP | CP001642 | Functions | AL123456 |
| pbpA/Rv0016c<br>(Class B PBP2) | AEJ48854 (482aa), nt.<br>20117-21565 | Penicillin-binding-protein, FtsI, **PbpA**, **pbp2** | CCP42738 (491aa), nt.<br>18759-20234 © |
| PonA1/Rv0050c<br>(Class A PBP1A) | AEJ48885 (784aa), nt.<br>54858-57212 | MrcB, Peptidoglycan<br>transglycosylase- D,D-<br>transpeptidase, **Pbp1a** | CCP42772 (678aa),<br>nt.53663-55699 |
| **Pbp4a**/Rv3627c<br>(class C; DacB) | AEJ49667 (473aa), nt.<br>1006473-1007894 | Endopeptidase/ serine<br>hydrolase, **AmpH1** | CCP43655 (532aa), nt.<br>1010136-1011734; 59aa<br>extension at NH2 |
| **Pbp4**/Rv1367c<br>AmpC | AEJ50426 (401aa), nt.<br>1537208-1538413 | β-lactamase, **AmpH2** | CCP44126 (377aa), nt.<br>1539512-1540645 © |
| Rv1497, β-lactamase<br>**LipL** esterase | AEJ50221 (429aa), nt.<br>1687057-1688346 | β-lactamase, **AmpH3** | CCP44258 (429aa) nt.<br>1689230-1687944 © |
| blaDHA /Rv1730c<br>(AmpH) | Deletion (no protein), nt.<br>1945765-1947296 | CP000611/ABQ73489;<br>nt. 1957210-1958763;<br>(Mt H37Ra), **AmpH4** | CCP44496 (517aa), nt.<br>1955692-1957245 © |
| **blaC**/blaA/Rv2068c<br>(blaTEM) | (no protein), nt. 286834-<br>285911 +/- | CP000611/ABQ73843<br>nt. 2335786-2336709<br>(Mt H37Ra) **blaC** | CCP44842<br>(307aa); nt. 2325886-2326809 © |
| pbpB/Rv2163c<br>(Class B PBP2) | AEJ50811 (605aa), nt.<br>2417473-2419290 | FtsI domain, **PbpB**,<br>D,D- transpeptidase | CCP44940 (679aa), nt.<br>2425048-2427087 © |
| PbpC/Rv2864c<br>(MecA) | AEJ51432 (582aa), nt.<br>3156295-3158043 | FtsI domain,<br>transpeptidase<br>lipoprotein /**mec A_N** | CCP45666 (603aa)<br>nt. 3175454-3177265 © |
| DacB2/Rv2911c<br>(Class C PBP7) | AEJ51473 (287aa), nt.<br>3199285-3200160 | **DacB2**, D,D-carboxypeptidase | CCP45713 (291aa), nt.<br>3218339-3219214 |
| DacB1/ Rv3330c<br>(Class C PBP6) | AEJ51876 (392aa) nt.<br>3707205-3708387 | DacB1, D,D-carboxypeptidase | CCP46151 (405aa), nt.<br>3715777-3716994 |
| Rv3627c<br>Conserved<br>protein | AEJ52151 (450aa), nt.<br>4059924-4061309<br>V2S mutation | DD-carboxypeptidase;<br>**PBP4** | CCP46450 (461); nt.<br>4065900-4067284 |
| PonA2/Rv3682<br>(Class A Pbp1B) | AEJ52200 (810aa), nt.<br>4116358-4118790 | MrcB, **PonA2**, D,D-transglycosylase-<br>transpeptidase, **Pbp1b** | CCP46506 (810aa), nt.<br>4121916-4124348 |
| Rv3762c<br>β-lactamase? | AEJ52278 (626aa), nt.<br>4201440-4203320 | Alkyl sulfatase and<br>MBL-hydrolase? | CCP46589 (626aa), nt.<br>4206996-4208876 © |
| Note: Rv0907 located at nt. 1687060-1688346 or NP_215422 (532aa) same to CCP43655 or AmpH.<br>Rv1922 is a lipoprotein (CCP44689; 371aa) and may not be PBP. |  |  |  |

Then, we organized many *Mt* low molecular weights class-C PBPs in a table as compared to *E. coli* class-C enzymes including blaTEM, blaAmpC and blaOXA-58 which was a signature of carbapenemase activity (table-3). The table stated that few PBPs designated as PBP4 but had no similarity to *E. coli* PBP4 suggesting wrong nomenclature (red). It was clearly found such enzymes had similarity to AmpH class-C enzymes and such information was wrong many sequences. We continued our search to *Mt* genomes and we had shown in table-4 that β- lactamases reported in *Mt* GG-36-11 (AN:CP025597) and *Mt* FDAARGOS_757 (AN: CP054013) as compared to *Mt* H37Rv genome (AN: AL123456): The data is important as we have identified the authentic PBP4 in *Mt* strain FDAARGOS_757 (protein id. AUP69687) with 34.5% similarity to *E coli* PBP4 (table-3). BLASTP search identified numerous *Mt* PBP4 but data still disclosed as conserved protein, unidentified protein or simple β-lactamase-like protein. It was thus clear that *Mt* PBP4 was wrongly described in many papers. Likely, such information was known by some scientists and as we saw that in *Mt* FDAARGOS_757 chromosome (AN:CP054013) there was no mention of any PBPs like PBP1, PBP2, PBP4 or PBP5 but disclosed as β-lactamase. Thus, PonA1 or PonA2 (class-A) or PBPA or PBPB (class-B) nomenclature are very popular and DacB (PBP5), DacB1 (PBP6) and DacB2 (PBP7) nomenclature are also getting easier to describe *Mt* PBPs. However, concept of *Mt* PBP3 and PBP4 are quite misleading everywhere in the database. Because PBPA and PBPB of *Mt* had molecular weight difference, some scientist described as PBP2 and PBP3. In the table-3 we showed many PBP4 described were truly had no similarity to *E. coli* PBP4 and they were blaAmpC class enzymes (upper panels). Table. 2A suggested PBPA and PBPB had blaOXA- 23 activities whereas PonA1 and PonA2 had no similarity to any β-lactamases. table-2B however, disclosed that DacB1(PBP6) and DacB2 (PBP7) had blaTEM similarities whereas very important and potent similarities of AmpH1, AmpH2, AmpH3 and AmpH4 enzymes with all class-C β-lactamases like blaAmpC, blaACC, blaDHA as well as blaCMY to lesser extent. It was known that class-C β-lactamases control the hydrolysis of cephalosporin drugs but not carbapenem drugs. The multi-alignment similarity of those AmpH isomers were shown in figure-2. It was found that the similarities of AmpH1, AmpH2, AmpH3 and AmpH4 proteins with *E. coli* AmpH protein obtained as 22.6% similarity/87% cover, 26.55% similarity/49% cover, 28.87% similarity/49% cover and 23.81% similarity/88% cover respectively but the low similarity for AmpH2 and AmpH3 as % cover was found. Still consistence similarity with blaAmpC were very important for blaAmpH1, blaAmpH2 and blaAmpH4 but the blaAmpH3 had no similarity to blaAmpC but to blaDHA (23.5%/26% cover). Importantly, *Mycobacterium* AmpH proteins were conserved among species and their AmpC similarities were described in the database. Thus, we are very optimistic about their functions as class-C β-lactamases. Because the BLAST-2 alignment of a plasmid AmpH protein (protein id. WJR74365) from *Enterobacter hormaechei* shared 23.2%/47%, 26.71%/67%, 30%/10% and 24.61%/79% (similarity/cover) with AmpH1, AmpH2, AmpH3 and AmpH4 enzymes of *Mt* respectively. Thus, the homology cover increased for blaAmpH2 but further decreased for blaAmpH3 of *Mt*. But when we did similar search with a *Vibrio parahaemolyticus* chromosomal AmpH protein (protein id. KKY41027), we found a decent homology 29.95%/ 49% cover for blaAmpH3 of *Mt* suggesting we were very right in our analysis of class-C β-lactamases. Similarly, we also BLAST-2 analysed with *Yersinia pekkanenii* AmpH (protein id. CNI55383) to get good homologies with *Mt* AmpH proteins as follows: AmpH1=24.05%/56%; AmpH2=27.85%/45%; AmpH3=25.77%/45% and AmpH4=24.13%/71% (% similarity/ % cover) and very similar as compared with *E. coli* AmpH enzyme (see, S9).

**Fig. 2.**
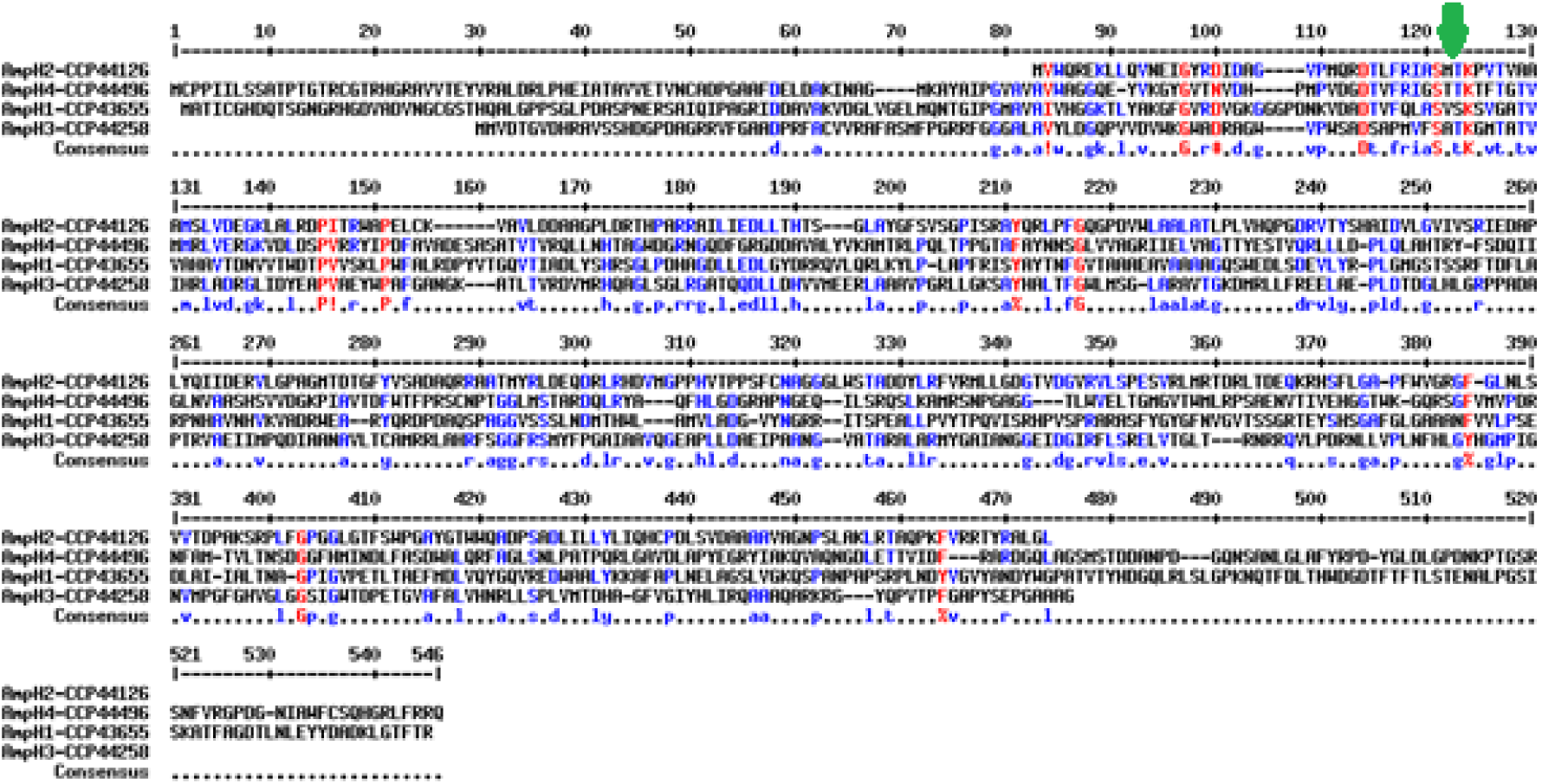
Demonstration of conserved SxxK motif in AmpH1, AmpH2, AmpH3 and AmpH4 *M. tuberculosis* AmpC class-C β-lactamases with 23-25% homology and 61-91% cover.

**Table-2.**
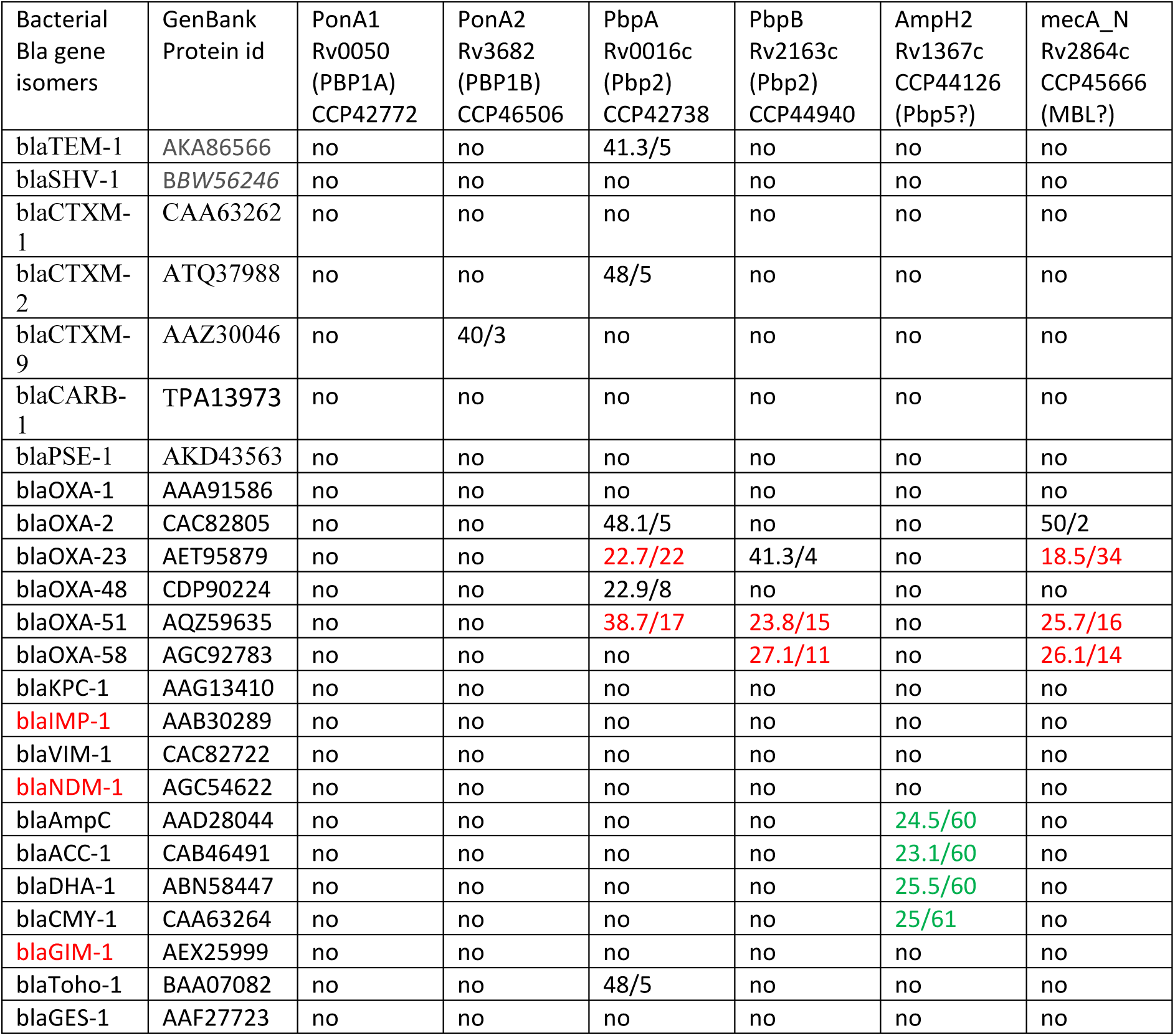
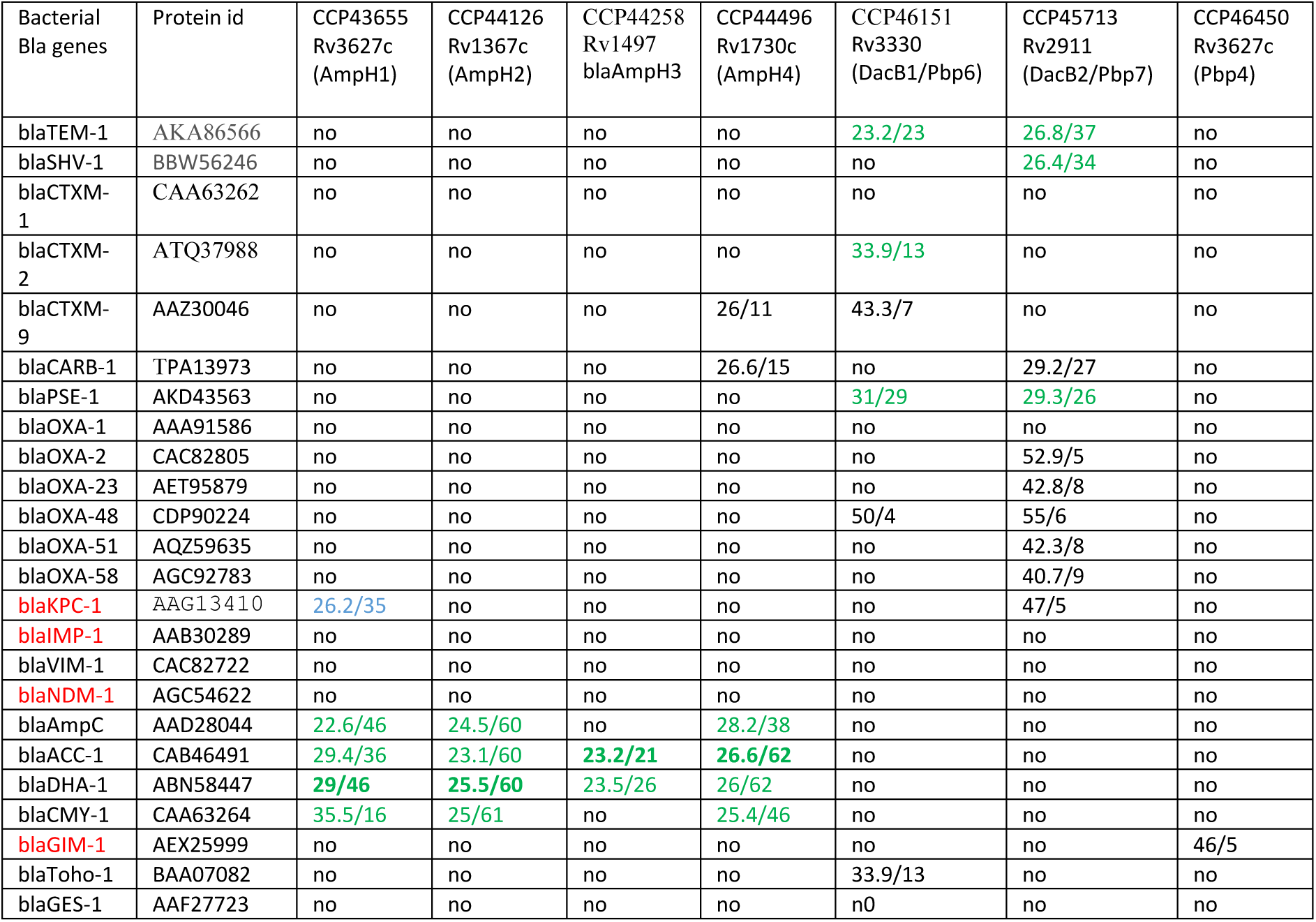
A: Demonstration of different *M. tuberculosis* class-A (PonA1 and PonA2) and class-B (PbpA and PbpB) PBPs homologies with different plasmid-mediated β-lactamases B: Demonstration of different Class-C β-lactamases in *Mycobacterium tuberculosis* (% similarity/ % cover); (no means no similarity at all during BLAST-2 search)

| Table-2A: Demonstration of different *M. tuberculosis* class-A (PonA1 and PonA2) and class-B (PbpA and PbpB) PBPs homologies with different plasmid-mediated β-lactamases |  |  |  |  |  |  |  |
| --- | --- | --- | --- | --- | --- | --- | --- |
| Bacterial<br>Bla gene<br>isomers | GenBank<br>Protein id | PonA1<br>Rv0050<br>(PBP1A)<br>CCP42772 | PonA2<br>Rv3682<br>(PBP1B)<br>CCP46506 | PbpA<br>Rv0016c<br>(Pbp2)<br>CCP42738 | PbpB<br>Rv2163c<br>(Pbp2)<br>CCP44940 | AmpH2<br>Rv1367c<br>CCP44126<br>(Pbp5?) | mecA_N<br>Rv2864c<br>CCP45666<br>(MBL?) |
| blaTEM-1 | AKA86566 | no | no | 41.3/5 | no | no | no |
| blaSHV-1 | BBW56246 | no | no | no | no | no | no |
| blaCTXM-1 | CAA63262 | no | no | no | no | no | no |
| blaCTXM-2 | ATQ37988 | no | no | 48/5 | no | no | no |
| blaCTXM-9 | AAZ30046 | no | 40/3 | no | no | no | no |
| blaCARB-1 | TPA13973 | no | no | no | no | no | no |
| blaPSE-1 | AKD43563 | no | no | no | no | no | no |
| blaOXA-1 | AAA91586 | no | no | no | no | no | no |
| blaOXA-2 | CAC82805 | no | no | 48.1/5 | no | no | 50/2 |
| blaOXA-23 | AET95879 | no | no | 22.7/22 | 41.3/4 | no | 18.5/34 |
| blaOXA-48 | CDP90224 | no | no | 22.9/8 | no | no | no |
| blaOXA-51 | AQZ59635 | no | no | 38.7/17 | 23.8/15 | no | 25.7/16 |
| blaOXA-58 | AGC92783 | no | no | no | 27.1/11 | no | 26.1/14 |
| blaKPC-1 | AAG13410 | no | no | no | no | no | no |
| blaIMP-1 | AAB30289 | no | no | no | no | no | no |
| blaVIM-1 | CAC82722 | no | no | no | no | no | no |
| blaNDM-1 | AGC54622 | no | no | no | no | no | no |
| blaAmpC | AAD28044 | no | no | no | no | 24.5/60 | no |
| blaACC-1 | CAB46491 | no | no | no | no | 23.1/60 | no |
| blaDHA-1 | ABN58447 | no | no | no | no | 25.5/60 | no |
| blaCMY-1 | CAA63264 | no | no | no | no | 25/61 | no |
| blaGIM-1 | AEX25999 | no | no | no | no | no | no |
| blaToho-1 | BAA07082 | no | no | 48/5 | no | no | no |
| blaGES-1 | AAF27723 | no | no | no | no | no | no |

| Table-2B: Demonstration of different Class-C β-lactamases in *Mycobacterium tuberculosis* (% similarity/ % cover); (no means no similarity at all during BLAST-2 search) | | | | | | | | |
| --- | --- | --- | --- | --- | --- | --- | --- | --- |
| Bacterial Bla genes | Protein id | CCP43655<br>Rv3627c<br>(AmpH1) | CCP44126<br>Rv1367c<br>(AmpH2) | CCP44258<br>Rv1497<br>blaAmpH3 | CCP44496<br>Rv1730c<br>(AmpH4) | CCP46151<br>Rv3330<br>(DacB1/Pbp6) | CCP45713<br>Rv2911<br>(DacB2/Pbp7) | CCP46450<br>Rv3627c<br>(Pbp4) |
| blaTEM-1 | AKA86566 | no | no | no | no | 23.2/23 | 26.8/37 | no |
| blaSHV-1 | BBW56246 | no | no | no | no | no | 26.4/34 | no |
| blaCTXM-1 | CAA63262 | no | no | no | no | no | no | no |
| blaCTXM-2 | ATQ37988 | no | no | no | no | 33.9/13 | no | no |
| blaCTXM-9 | AAZ30046 | no | no | no | 26/11 | 43.3/7 | no | no |
| blaCARB-1 | TPA13973 | no | no | no | 26.6/15 | no | 29.2/27 | no |
| blaPSE-1 | AKD43563 | no | no | no | no | 31/29 | 29.3/26 | no |
| blaOXA-1 | AAA91586 | no | no | no | no | no | no | no |
| blaOXA-2 | CAC82805 | no | no | no | no | no | 52.9/5 | no |
| blaOXA-23 | AET95879 | no | no | no | no | no | 42.8/8 | no |
| blaOXA-48 | CDP90224 | no | no | no | no | 50/4 | 55/6 | no |
| blaOXA-51 | AQZ59635 | no | no | no | no | no | 42.3/8 | no |
| blaOXA-58 | AGC92783 | no | no | no | no | no | 40.7/9 | no |
| blaKPC-1 | AAG13410 | 26.2/35 | no | no | no | no | 47/5 | no |
| blaIMP-1 | AAB30289 | no | no | no | no | no | no | no |
| blaVIM-1 | CAC82722 | no | no | no | no | no | no | no |
| blaNDM-1 | AGC54622 | no | no | no | no | no | no | no |
| blaAmpC | AAD28044 | 22.6/46 | 24.5/60 | no | 28.2/38 | no | no | no |
| blaACC-1 | CAB46491 | 29.4/36 | 23.1/60 | 23.2/21 | 26.6/62 | no | no | no |
| blaDHA-1 | ABN58447 | 29/46 | 25.5/60 | 23.5/26 | 26/62 | no | no | no |
| blaCMY-1 | CAA63264 | 35.5/16 | 25/61 | no | 25.4/46 | no | no | no |
| blaGIM-1 | AEX25999 | no | no | no | no | no | no | 46/5 |
| blaToho-1 | BAA07082 | no | no | no | no | 33.9/13 | no | no |
| blaGES-1 | AAF27723 | no | no | no | no | no | no | no |

**Table-3:**
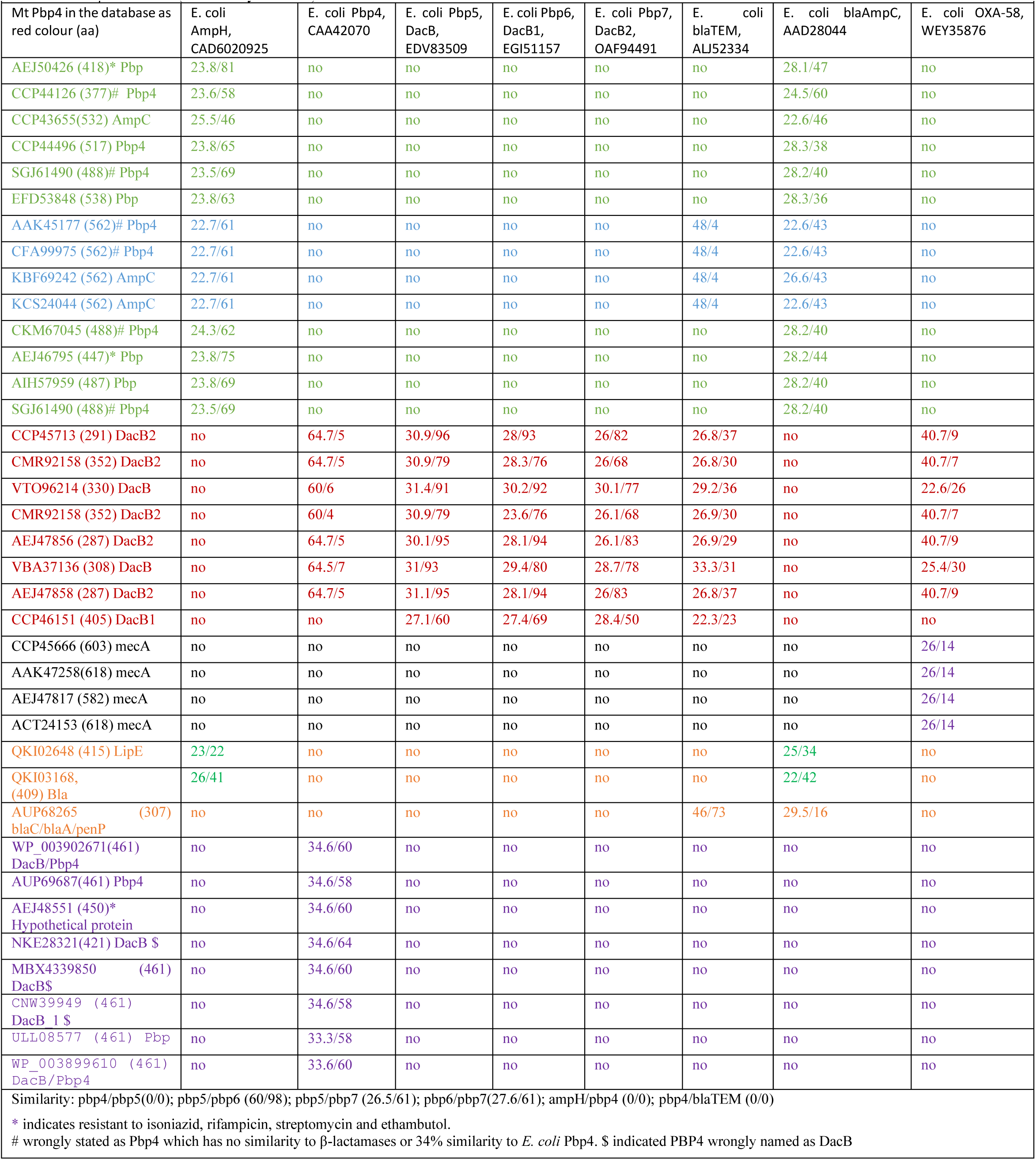
The h`omology among` E`. coli`different *Mt* PBPs described as class C penicillin-binding-protein (Pbp4 in majority cases) in the database as compared to *E. coli* PBPs and β-lactamases (% similarity/ % cover)

**Table-4:** Demonstration of PBPs and Beta-lactamases reported in *Mt* GG-36-11 (AN: CP025597) and *Mt* FDAARGOS_757 (AN: CP054013) as compared to *Mt* H37Rv genome (AN: AL123456): This data is important as identified authentic PBP4 in *Mt* strain FDAARGOS_757 (+1 site in the chromosomes vary).

| Name of the PBP in *Mt* GG-36-11 | Protein id for *Mt* GG-36-11 | AL123456 homology | Name of the PBP in FDAARGOS_757 | Protein id for FDAARGOS_757 | AL123456 homology % |
| --- | --- | --- | --- | --- | --- |
| PbpA | AUP66337 | CCP42738 (100/100) **Pbp2a** | DacB1 carboxypeptidase | QKI02248 | CCP46151 (100/100) **DacB1** |
| PBP1A | AUP66368 | CCP42772 (99.7/82) **PonA1** | PonA2 Transglycosylase/ D,D-transpeptidase | QKI02574 | CCP46506 (100/100) **PonA2 (PBP1B)** |
| AmpC | AUP70055 | CCP43655 (98.7/100) **AmpH1?** | D-Alanyl-D-Ala-Carboxypeptidase , DacB | QKI02525 | CCP46450 (100/100) **Pbp4** |
| β-lactamase | AUP67611 | CCP44126 (100/100) **AmpH2** | Beta-lactamase family protein (lipE) | QKI02648 (lipase, LipE) **New-1** | CCP43655 (24.2/57) |
| β-lactamase | AUP67734 | CCP44258 (100/100) **AmpH3** | PbpA D,D-transpeptidase | QKI02805 | CCP42738 (100/100) **Pbp2 (Fts1)** |
| PBP | AUP67942 | CCP44496 (100/100) **AmpH4** | PonA1 Transglycosylase-DD-transpeptidase | QKI02837 | CCP42772 (99.5/100) **PonA1(PBP1A)** |
| **blaA** / blaC | AUP68265 | CCP44842 (100/100) **blaC** | Beta-lactamase family protein | QKI03645 (Serine hydrolase) | CCP43655 (99.6/96) **AmpH1?** |
| PbpB | AUP68350 | CCP44940 (100/100) **Pbp2b** | Beta-lactamase family protein | QKI04082 | CCP44126 (100/100) **AmpH2** |
| PBP | AUP68992 | CCP45666 (100/100) **MecA_N** | Beta-lactamase | QKI04208 | CCP44258 (100/100) **AmpH3** |
| D-alanyl-D-alanine carboxypeptidase | AUP69035 | CCP45713 (100/100) **DacB2** | Beta-lactamase family protein | QKI04417 | CCP44496 (100/100) **AmpH4** |
| D-alanyl-D-alanine carboxypeptidase | AUP69421 | CCP46151 (100/100) **DacB1** | blaA /penP | QKI04734 | CCP44842 (100/100) **blaC** |
| D,D-endopeptidase | AUP69687 | CCP46450 (100/100) **PBP4** | PbpB D,D-transpeptidase | QKI04820 | CCP44940 (100/100) **PbpB** |
| Membrane carboxypeptidase | AUP69732 | CCP46506 (100/1000) **ponA2** | Beta-lactamase | QKI05481 | CCP45666 (100/100) **MecA_N** |

Note: AUP66689 homologues: AIH44463, AGL25868, KBI52351, AFY28928

The interesting finding that many AmpH enzymes were described as Pbp4 as seen in protein ids. CNW35153, CKM56251, CMR84911, CMA68255, CKR07362, CFS29512, COW38226 and CKR60184 (see, table-3). The serine hydrolases (protein ids. OBA76072, OBA64464, KUP04565) had very strong similarities to AmpH enzymes (protein ids. VBA58402, VBA42298) and almost indistinguishable by BLASTP search. Similarly, lipL esterase enzyme described in the *Mt* chromosome database also had strong similarity to *E. coli* AmpH β- lactamase. Combined together, we selected blaAmpH1, blaAmpH2, blaAmpH3 and blaAmpH4 as true class-C β-lactamases of *M. tuberculosis* controlling hydrolysis of cephalosporin drugs. The BLAST-2 similarities were shown in figure-3 and figure-4 demonstrating quite 30% homology between them. Although, multiple alignment reduced such similarity (figure-2).

**Fig. 3.**
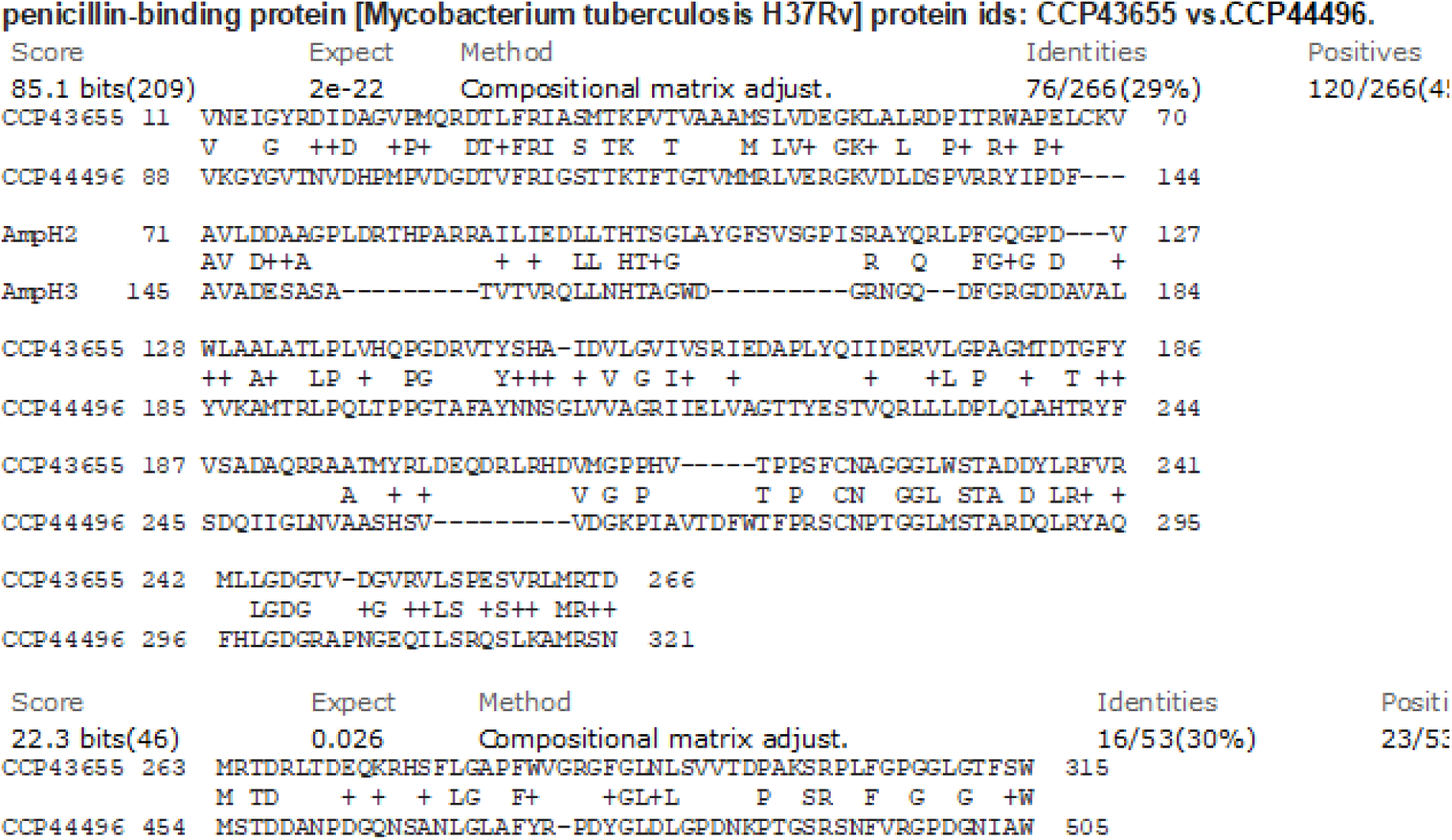
BLAST-2 similarity between AmpH2 and AmpH3 β-lactamases of *M. tuberculosis*.

**Fig. 4.**
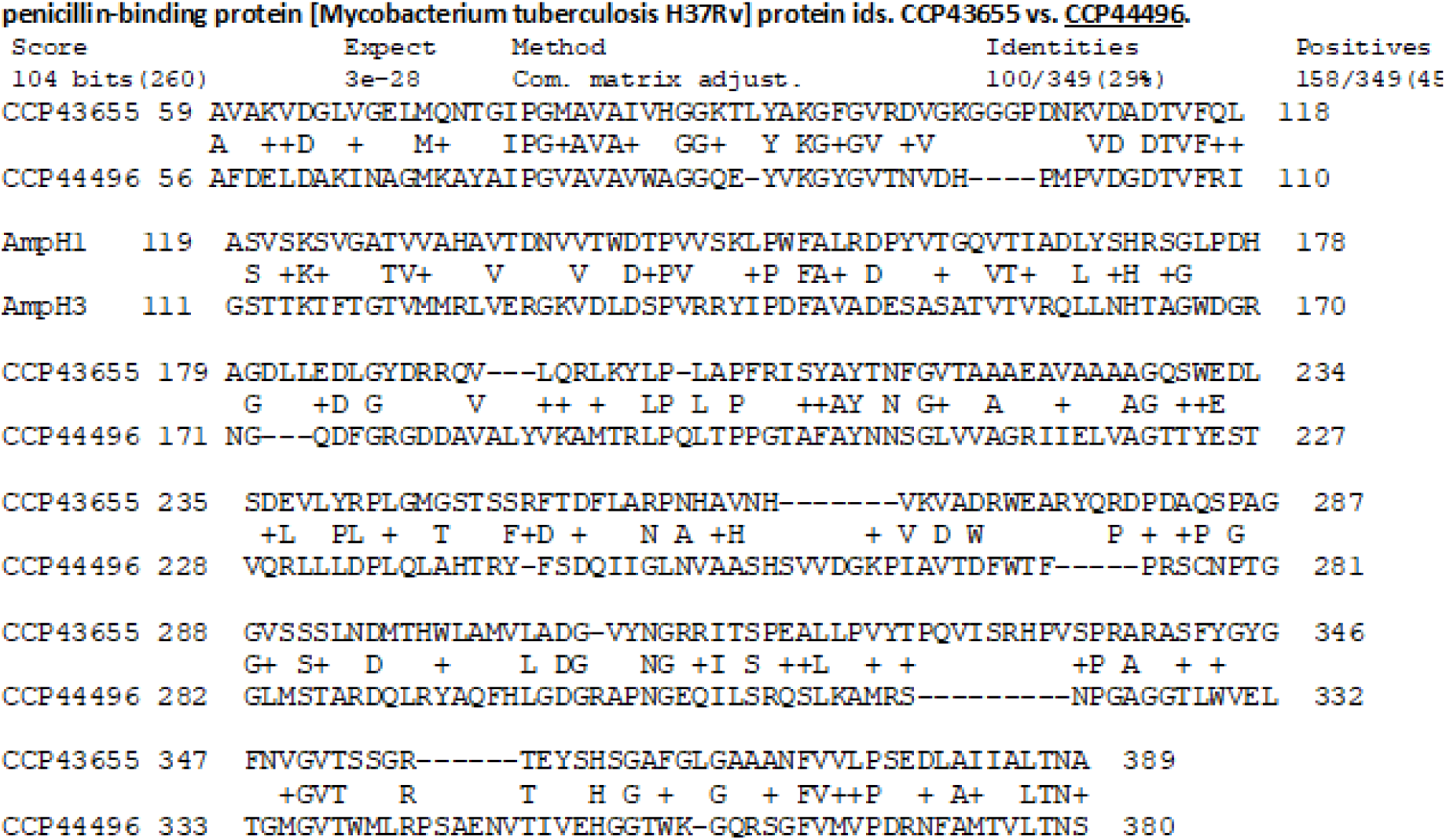
BLAST-2 similarity between AmpH1 and AmpH3 PBPs of *M. tuberculosis*.

**Fig. 5.**
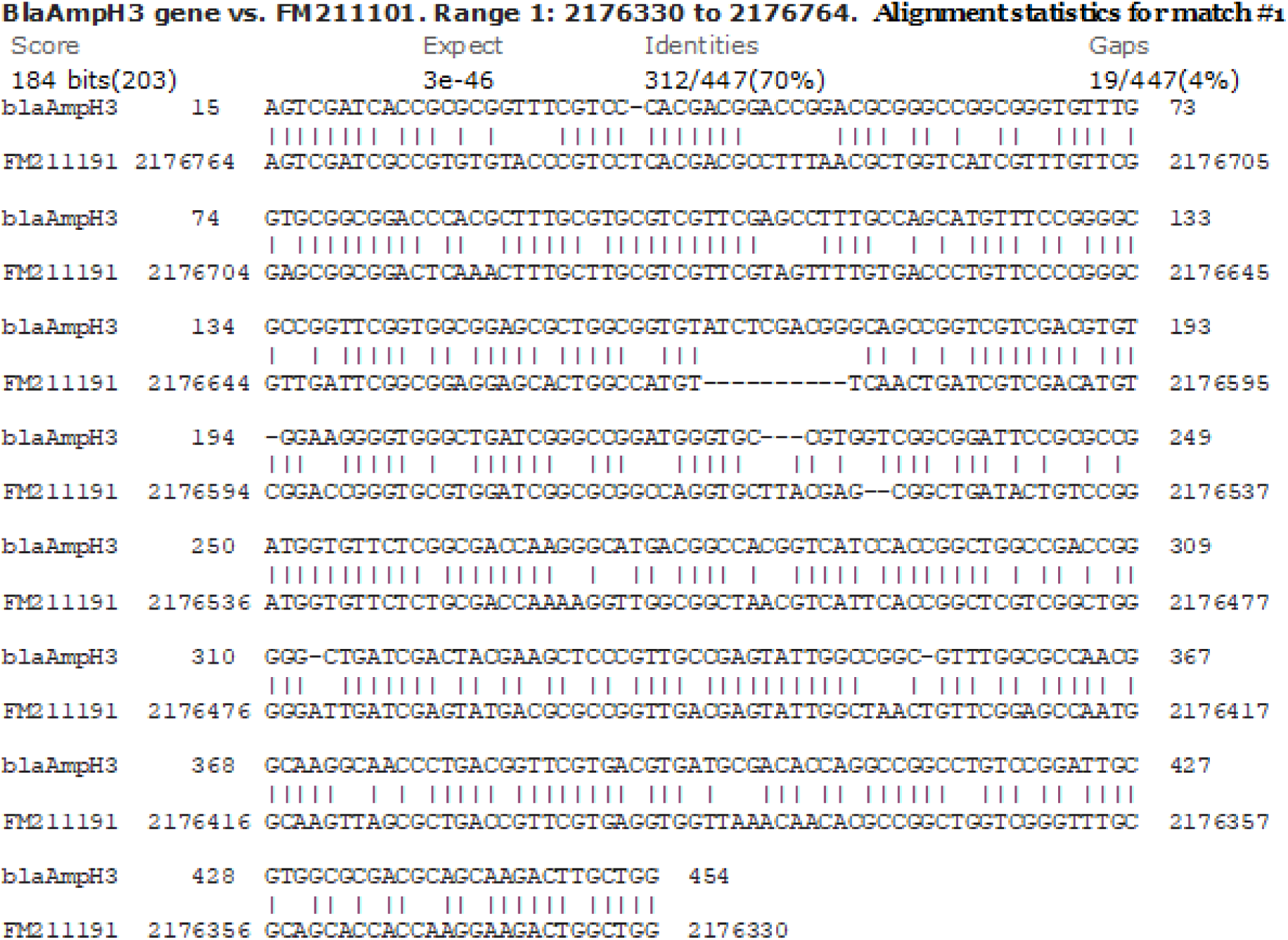
BLASTN similarity between blaAmpH3 gene and *M. leprae* genome (match-1 called lipL esterase pseudogene).

**Fig. 6.**
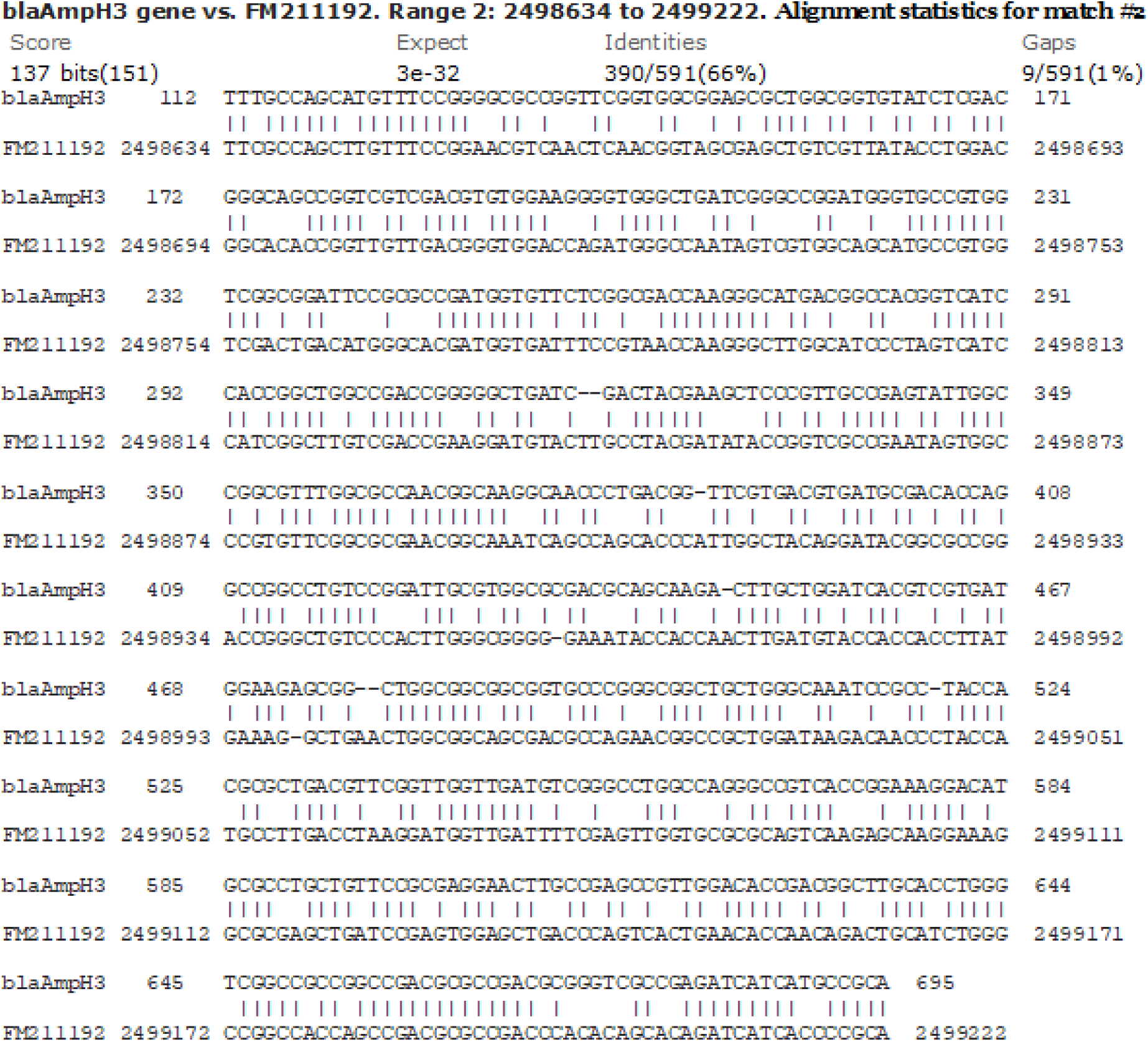
BLASTN similarity between blaAmpH3 gene and *M. leprae* genome (match-2 also called lipL esterase pseudogene). The blaAmpH1 gene similarity located at nt. 2516623 and blaAmpH2 gene similarity located in nt. 642079 of the *M. leprae* genome (acc. nos. FM211192, CP029543 and AP014567). But no such similarity for blaAmpH4 gene (data not shown).

**Fig. 7.**
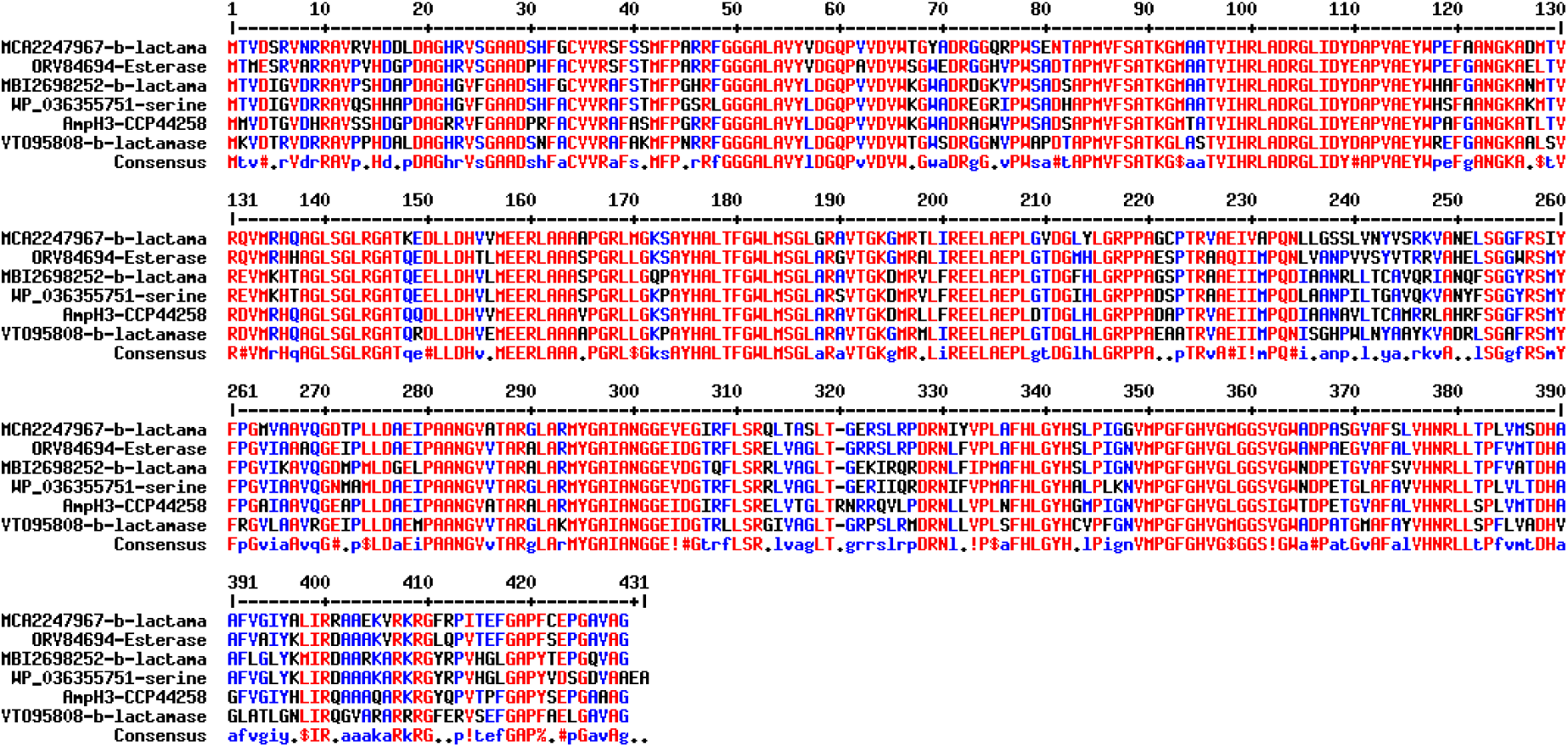
Demonstration of blaAmpH3 homologue in *M. intracellulare* (JAEKMP010000010), *M. asiaticum* (WP_036355751), *M. interjectum* (LQPD01000066) and other *M. species* (JACPNU010000009). The data indicated that blaAmpH3 class-c enzymes of *Mt* has strong similarity to esterase, serine hydrolase and other β-lactamases reported in the database. Actually, such enzymes were AmpH3 class and has less similarity to blaAmpH1, blaAmpH2 and blaAmpH4.

We compared the *M. tuberculosis* PBPs with *M. leprae* enzymes (see, S4) because blaAmpH1/2/3 were detected through BLAST-2 search with chromosomes of *M. leprae* except blaAmpH4. But scanning of WGS were unable to detect the AmpH protein at the chromosomal locus and only found written as pseudogene. The two sequences obtained from homology search with *Mt* blaAmpH3 gene produced two homology positions. Such two homology sequences when BLAST-2 compared gave homology portions with huge gap (figure-8). The BLASTX search of AmpH1.1 fragment produced homology peptides but one small peptide with *M. leprae* and many with other *Mycobacterium* species including *M. tuberculosis*. Multi-alignment of those protein clearly indicated such proteins were AmpH3 class as expected but had some mutations among them as in case of an esterase gene or a lipoprotein called lipL gene (figure-9). Similarly, repeated experiment with AmpH1.1 and AmpH1.2 fragments of *M. leprae* produced some homology but no great insertion/deletion was found (one or two nucleotides only). The BLASTX homology detected many protein sequences described as Pbp4 of *M. tuberculosis* but as expected were blaAmpH1 enzymes with some difference at the NH2 terminus initiation point (figure-10). Interestingly, again we got two small peptide homologies (91 aa and 81 aa long) in *M. leprae* pinpointing the pseudogene concept. However, we detected PonA1 (protein id. CAC32220), PonA2 (protein id. CAC31824), PBPA (protein id. AWV47033), PBPB (protein id. AWV47654), MecA_N (protein id. AWV48102), DacC (protein id. AWV47526), DacB1(protein id. AWV47188) all PBPs except blaAmpH1/2/3/4 enzymes. Rationally, *M. tuberculosis* genome is 4.46 million bases whereas *M. leprae* has only 3.25 million bases. We speculated that deletions and insertions including point mutations were occurred at the AmpH locus of *M. leprae* [52]. Thus, we explained the AmpH pseudogenes concept but needed more careful analysis. Never-the-less, we beautifully disclosed the present concept of PBP4 which in most cases were AmpH genes of *Mt* and true PBP4 was never analysed in *Mt*. The *AmpH* gene of many bacteria shared close similarities as demonstrated in figure-11. But many mutations in the *V. parahaemolyticus* chromosomal AmpH protein (protein id. KKY4107), *Enterobacter hormaechei* plasmid-mediated AmpH protein (protein id. WJR74365) and *Raoviltella planticola* AmpH protein (protein id. UNK76796) were found. Truly, the database also pointed penicillin-binding-protein 4, has AmpC motif which was not possible pinpointing wrong nomenclature (table-3, upper panels). That way we got strength of idea that PBP4 ideally first enzyme of class-C PBP and scientist sometime became over smart to describe the fact and assumed all class-C enzymes as PBP4. As for example, PBP2B sometimes described as PBP3 in *M. tuberculosis* and an AmpH gene disclosed as PBP5. Ideally, *E. coli* PBP3 and PBP5 had no homologies to *M. tuberculosis* counterpart and such misleading nomenclature appeared in the PubMed and GenBank. In table-3 we also shown few PBP4 which appeared as AmpC class enzymes with similarities to blaAmpC, blaDHA, blaACC and blaCMY that all smartly hydrolysed the cephalosporins like cefotaxime which widely used now to clear infections. We have not found good similarity to MBLs and carbapenemases (blaNDM-1, blaVIM-1, blaIMP-1, blaOXA-23, blaOXA-58) with *M. tuberculosis* PBPs (table- 2). Whereas PonA1 and PonA2 disclosed as strong hydrolysing activities for carbapenem drugs and we opposed such act. Truly, we found some similarities of PBPA and PBPB (PBP2 class) with blaOXA-23 or we also found some similarity of blaMecA_N with blaOXA-58 enzyme and that way such enzyme might perform carbapenem binding and hydrolysis. In fact, blaC β- lactamase which also referred as blaA in some cases hydrolysed the penicillin drugs better whereas DacA1(Pbp6) and DacB2 (Pbp7) might be considered for such activities as demonstrated by homology search with 24 plasmid-mediated β-lactamases of *E. coli* and *K. pneumoniae* (table-3). The blaC enzyme of *M. tuberculosis* had similarity to blaZ of *S. aureus* and blaCARB of *V. parahaemolyticus* suggesting an important enzyme for penicillin hydrolysis. However, blaC also has similarities to blaCTX-M1/2/9 considering its activity as cephalosporinase (ESBL) [1]. Chromosomal enzymes were found inducible as in blaZ β- lactamase of *S. aureus* and AmpC enzyme of *S. marcescens* and *P. aeruginosa.* However, such induction was lost (blaR1 or ampR) in case such β-lactamase appeared in plasmid and likely multi-copies plasmids increased the gene dosage. Sadly, we are unable to show any induction system in *M. tuberculosis* and much careful attention may be necessary.

**Fig. 8.**
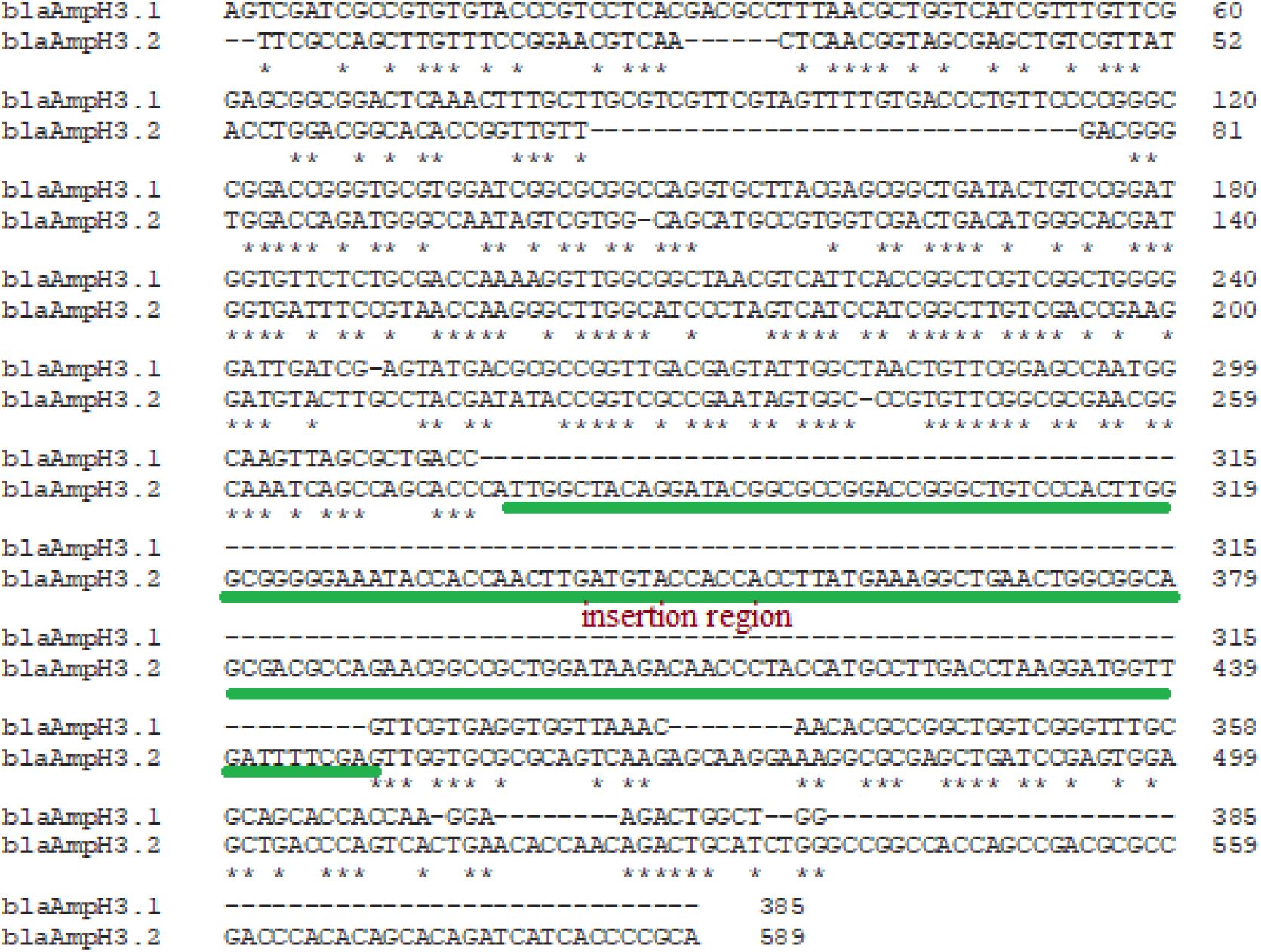
Demonstration of AmpH3 gene mutation, deletion and insertion to form a pseudogene in *M. leprae* genome. We found two regions homologies by BLAST search in *M. leprae* genome and such two regions were compared here to show a big insertion in second homology region of *M. leprae* genome producing a pseudogene of blaAmpH3 gene. Such insertion sequence did not produce any protein as determined by BlastX search but expected as an IS- element.

**Fig. 9.**
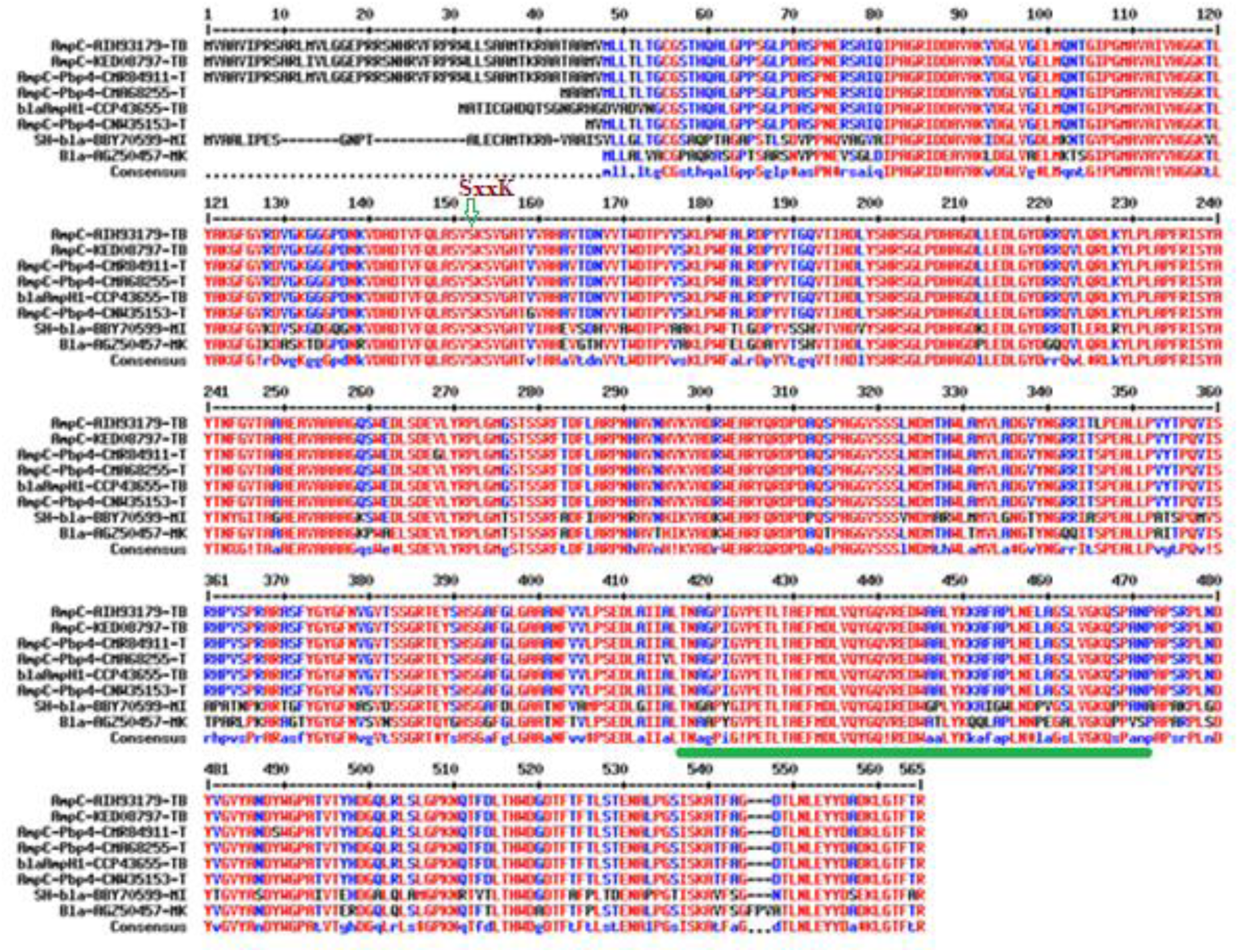
Demonstration of blaAmpH1 in *M. tuberculosis*, *M. kamsasii, M. intracellulare* described as Pbp4 as obtained by BLASTX search of *M. leprae* AmpH1.1 pseudogene fragment. In *M. leprae* two small peptides (71 and 91aa long, green underlined) with 80% similarities (protein ids. WP_239494153, WP_041323163) may be produced instead full- length protein. The NH2-terminal variations were found likely due to different GUG and UUG initiation codons usage over AUG universal codon. Some *Mt* Serine hydrolase (SH) has very similar sequence to AmpH proteins.

**Fig. 10.**
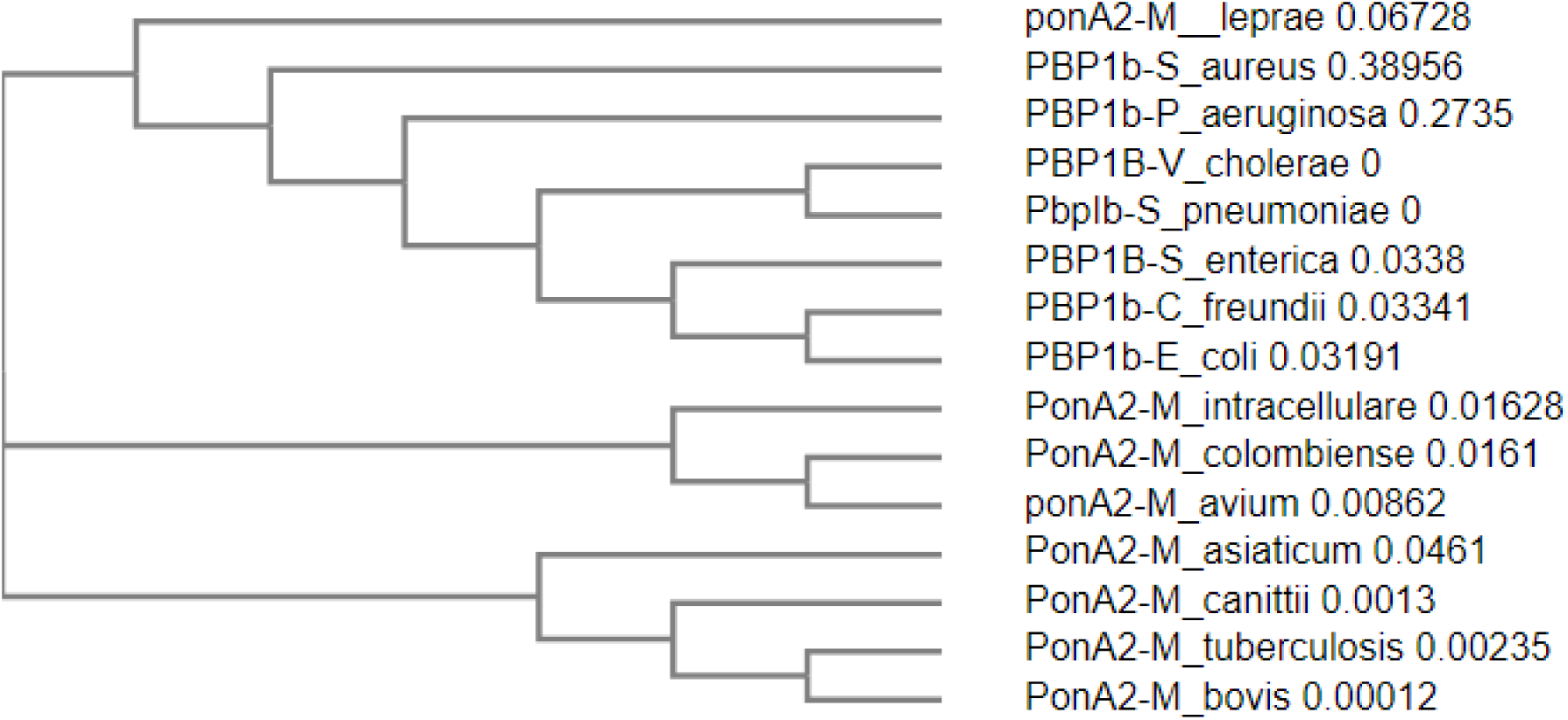
Phylogenetic relation of *M. leprae* ponA2 with other *Mycobacterium* species and common bacterial PonA2 proteins and surprisingly very closer to *S. aureus* PonA2. This data suggested that *M. leprae* evolution was quite complex and different from *M. tuberculosis*,*M. bovis* and *M. intracellulare* or *M. avium*. Note that PonA1, PonA2, PbpA, PbpB and MecA_N of *M. tuberculosis* and *M. leprae* have great BLASTP similarities.

**Fig. 11.**
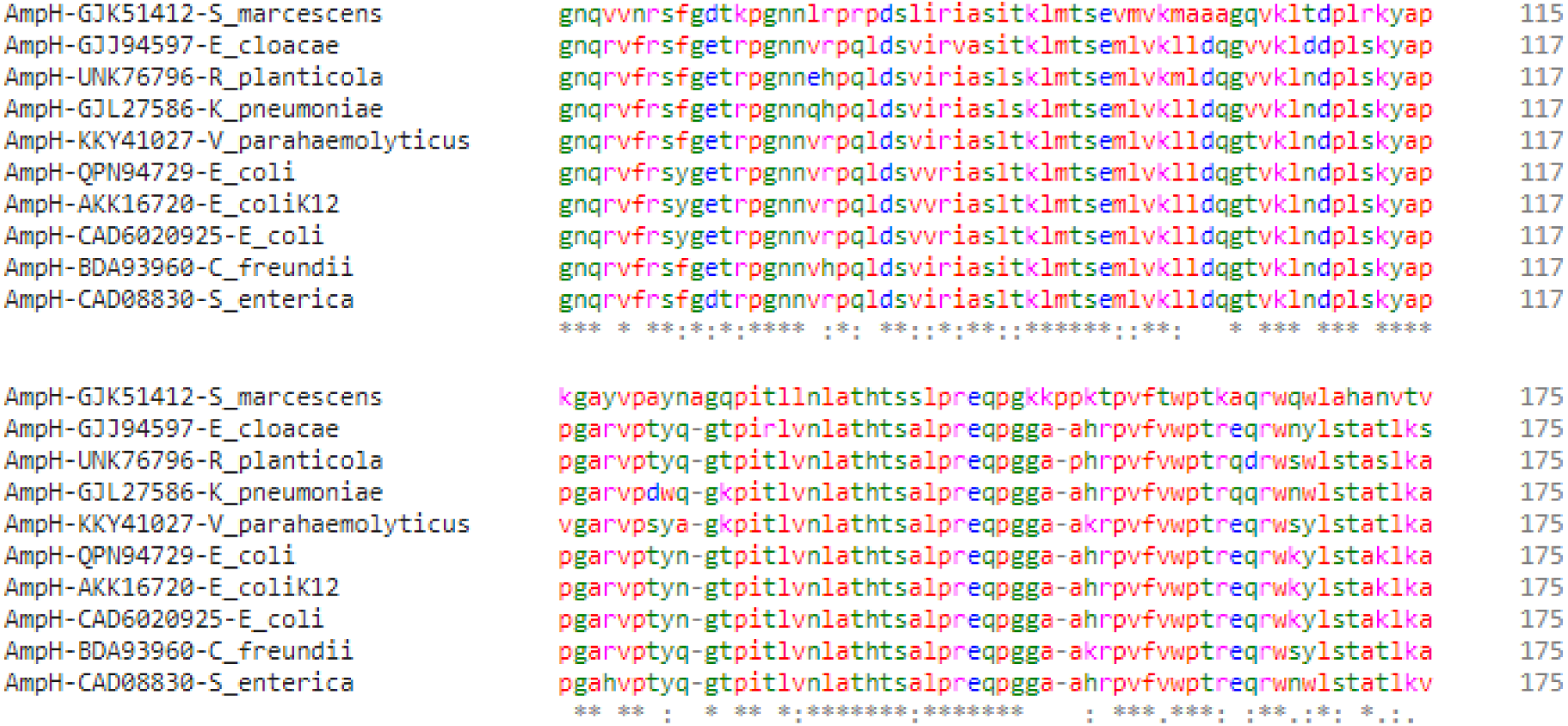
Carboxy-terminal homologies and mutations among AmpH proteins of bacteria (part of the alignment was shown). In *S. marcescens* one AA insertion (Alanine at 126 and Proline at 150 positions) was important including R59V, LLDQ=MAAA and A175V mutations.

However, sequence variation during sequencing possible and also SNP could be seen. Interestingly, in the *Mt* strain FDAARGOS_757, the AmpH1 gene SNP was found where one extra-G at nt70 of the AmpH1 gene creating a stop codon TGA at codon 45. However, from the upstream ATG codon the protein was produced with alter 22 AAs at the NH2-terminal. Astonishingly, we found same case in *Mt* strain GG-36-11 AmpH1 gene and it could a sequencing error. To clarify the issue, we BLASTN search 60nt with or without extra-G to see the penetration of the mutant allele. It appeared normal (*Mt* strain H37Rv) had only 46 sequences whereas rest thousand were with extra-G. However, such discrepancy did not alter the β-lactamase motif. Now question may appear if more AmpC-like genes in *M. tuberculosis*? Surely, we found a serine hydrolase protein in *Mt* strain FDAARGOS_757 with low similarity to AmpH1 (23%/42%) with protein id. QKI03168 implicated as β-lactamase as such domains spread to different esterase and lipoprotein and akyltransferases. More surprisingly, large plasmids (accession numbers: NC_012692, LN850163, LC055503, CP007558) with β- lactamases (blaTEM, blaCTXM, blaAmpC, blaSHV) did not show homology with *bla*AmpH1/2/3/4 genes but only 17nt homology (173580-5’-CCTTGGTGCCGTCGACC-3’- 173564) detected in the transposase (protein id. AHY14824) of *C. frundii* plasmid pKEC-a3c (AN:CP007558) with the blaAmpH4 gene nt. 1236-1252 of *M. tuberculosis*. Thus, all *bla*AmpH1/2/3/4 genes involve with higher rearrangement and mutations involving IS-elements. Interestingly, BLASTN search with this 17nt sequence selected many plasmids and chromosomes including many *M. tuberculosis* chromosomes. The *AmpC* or *AmpH* genes were chromosomal origin and their origin plasmids were further checked. Analysis found two *K. pneumoniae* plasmids (accession nos. CP067912 and AP024793) found homologies to 17nt sequence in the IS1182-like ISKpn6 transposase gene. Similarly, *C. freundii* plasmids (accession nos. CP054297 and CP110778) hybridised to 17nt sequence in the same IS1182- like ISKpn6 transposase gene. Such finding used to be crucial as we proposed that *bla*AmpH4 gene was produced by insertion of ISKpn6 transposase although BLASTP search produced no homology between blaAmpH4 protein and ISKpn6 transposase protein. The 17nt sequence however, detected blaAmpH4 sequence in the chromosome. The *Mt* strain 5521 (AN:CP127276, nt. 1946009-1945993) and *Mt* strain 02-R0894 (AN:CP089774, nt. 195118-1951102) selected by 17nt BLASTN search and in that locus blaAmpH4 protein sequence (protein ids.WJH80555 and WIY18944) were found (figure-12). However, we also selected a serine hydrolase with protein id. CKR72489, implicated as putative β-lactamase of *M. tuberculosis*. Thus, we confirmed the role of ISKpn6 sequence and a transposase in the genesis of *M. tuberculosis* conserved blaAmpH4 gene. Many β-lactamase proteins were selected and multi-alignment was done in presence of blaAmpH4 protein. The data presented in figure-12 that such β-lactamases were perfectly blaAmpH4 protein and only two AAs mutations (D435E and F495V) were found in the *Mt* strain 5521 (protein id. WIY18944) at the carboxy-terminus. We also detected a β-lactamase pseudogene in *Mt* FDAAGROS_756 that was occurred due to deletions of 19nt (5’-AACGGTCATCGACTTCCG-3’) at 1313nt and a 3nt (5’-CTT-3’) at 1481nt of AmpH4 gene (see, S5). Thus, roles of IS-elements (ISKpn6 transposase) in the creation of AmpH4 gene was postulated and similar events of deletions, mutations and rearrangements could be possible due to over dose use of penicillin, cephalosporin and carbapenem antibiotics.

**Fig. 12.**
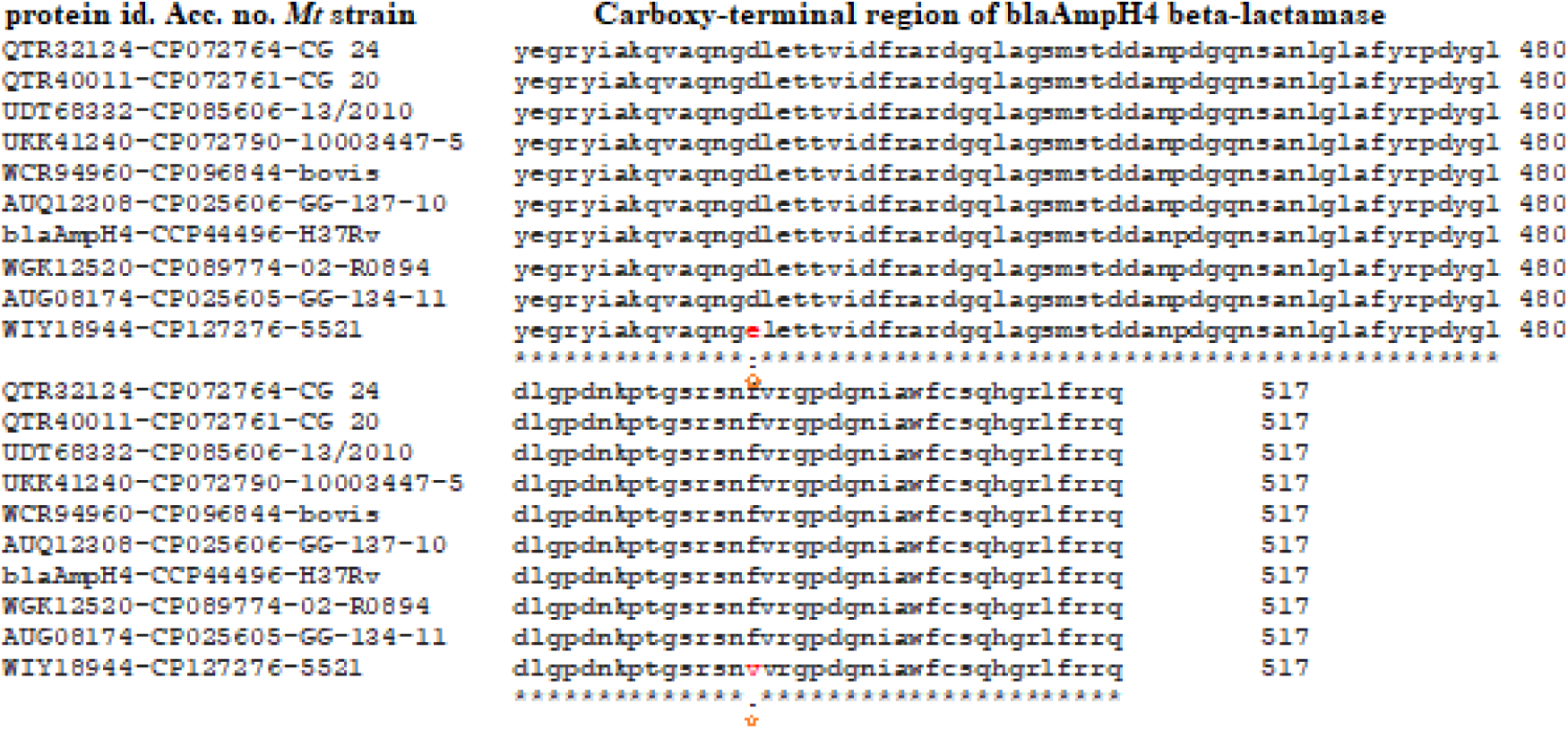
Demonstration of seventeen bases oligonucleotide (found in ISKpn6 transposase gene) BLASTN selected *Mt* chromosomes derived conserved blaAmpH4 proteins described as β- lactamase and only two mutations in ampH4 protein of *Mt* strain 5521 were detected (part of the alignment shown). The oligonucleotides only selected ampH4 protein and thus the generation of ampH1/2/3 may be done by other transposase and IS-elements.

We have decided to make PCR primers specific for all *Mt* PBPs for RT-PCR to detect their expression and also to check the individual enzyme localization after complete genome sequencing. Such attempt will surely avoid the case of nomenclature problem. The PonA1F primer is 5’-GTA CCA ACC AGG TCT CCA CG-3’ and PonA1R primer is 5’-TCG AGC AAC TCC CTT GTC AC-3’ with PCR product 718bp and Tm of the primers is 60°C. The PonA2F primer is 5’-CGA TTC CAG GCC AAA GGT CT-3’ and PonA2R primer is 5’-TCG ATG AGC TGG TCG ATT GG-3’ with PCR product 404bp and Tm 60°C. The PBPA primers were made and PbpAF= 5’-CTG CGG GTC TAT CCC AAT CC-3’ and PbpAR= 5’-CCG CCG TAG TTC TCT AGC TG-3’ with Tm=60^0^C and PCR product size=566bp. Similarly, the PBPB primers were made and PbpBF= 5’-CAC GCT GAT GCT TTC CCA AC-3’ and PbpBR= 5’-CCG GCG AAG GTG ATC CAA TA-3’ with Tm= 60^0^C and PCR product size= 516bp. The Pbp4F primers is 5’-AAC GCC TCC GAC AAT GTG AT-3’ and Pbp4R is 5’- AAC TTA GTG GCC AGA GCG TC-3’ with PCR product size 485bp and Tm for primers about 60°C. Both primers selected only single 100% match on the *Mt* genome: As for example, *Mt* H37Rv-1 genome selected nt. 4072802-4072821(+/-) for forward primer and nt. 4072337- 4072356 (+/+) for reverse primer (AN: CP110619) during BLASTN search of oligonucleotides separately.

The eight primers of blaAmpH1/2/3/4 isomers were also made and their localization in the *Mycobacterium* also confirmed and a restriction enzyme analysis scheme outlined (table-6 and table-7). It was found that restriction enzymes NotI (383, 173bp), NarI (349, 239bp), MscI (364, 181bp) and NdeI (307, 283bp) cut single in the AmpH1.1, ampH2.1, ampH3.1 and ampH4.1 PCR fragments respectively (table-7) allowing to differentiate the PCR fragments without DNA sequencing. The 1.2% agarose gel electrophoresis would be require to see the distinct low molecular weight bands in presence of ethidium bromide. Although the primers showed 100% homology with *Mt* H37Rv-1 and quite high similarities with *M. bovis* but *M. marinum, M. kansasii, M. avium, M. leprae* and *M. intracellulare* did not showed 100% homology. Thus, all primers could be assigned as *Mt* specific but high annealing temperature (54°C) should be maintained during PCR to reduce unwanted PCR bands.

**Table-6:**
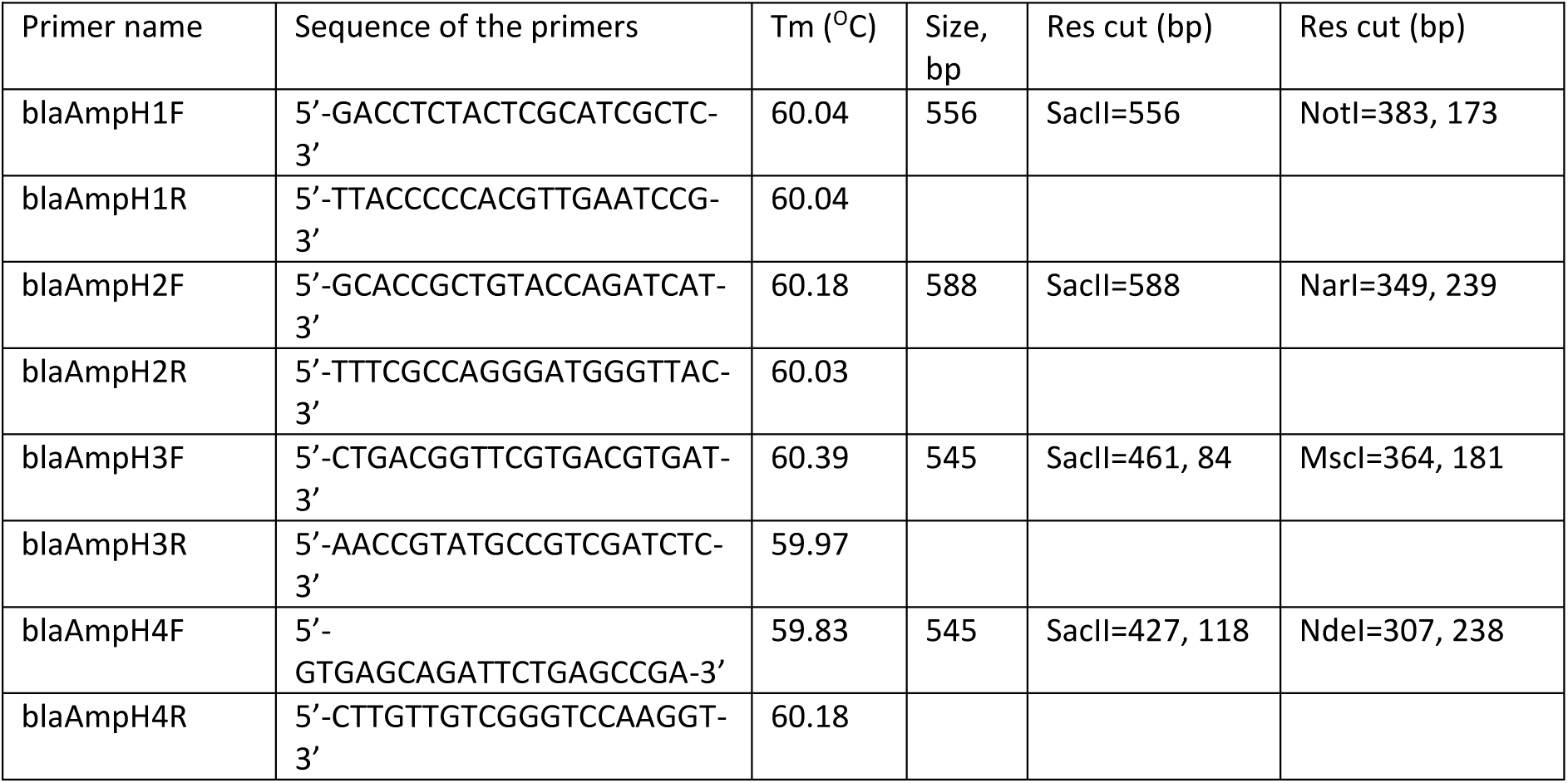
Primers for four BlaAmpH isomers and restriction enzyme analysis.

**Table-7:** Homology of AmpH1/2/3/4 primers with different *Mycobacterium* species.

| Primers name | *M. tuberculosis*<br>CP110619 | *M. bovis*<br>AP010918 | *M. avium*<br>CP033911 | *M. leprae*<br>AL450380 | *M. kansasii*<br>CP006835 | *M. intracellulare*<br>CP012885 |
| --- | --- | --- | --- | --- | --- | --- |
| blaAmpH1F | 1010638 | 1012826 | no | no | 2108763 | no |
| blaAmpH1R | 1011193 | 1013381 | no | no | no | no |
| blaAmpH2F | 1540162 | 1539557<br>A=G at 16* | no (8-19) | no (1-12) | no (3-14) | no (4-18) |
| blaAmpH2R | 1539575 | 1538970 | no (3-14) | no (3-15) | no (1-13) | no (2-14) |
| blaAmpH3F | 1688535 | 1686288 | no (5-17) | no (2-15) | no (3-16) | no (3-16) |
| blaAmpH3R | 1689079 | 1686832 | no (8-20) | no (7-19) | no (7-20) | no (8-20) |
| blaAmpH4F | 1956543 | 1934390 | no (3-14) | no (9-20) | no (1-12) | no (3-20)# |
| blaAmpH4R | 1955999 | 1933846 | no (7-20) | no (8-19) | 1665625<br>A=G at 16* | no (7-19) |

Note: the position in the genome with 100% similarity shown. In the bracket ( ) homology region in the 20nt primers were given. \* indicated one nucleotide mismatch only. # indicated quite high homology

The primers for mecA_N gene (protein id. CCP45666) were also made and mecAF= 5’-CAT CCC AAG CTA GGC GAA CA-3’ and mecAR= 5’-CAA CGT GAT CGA AAC CGA CG-3’ with Tm 60^0^C and the product size=522bp. The mecA_N has similarity to blaOXA-58 (19%/58% cover) and also BLASTP search identified conserved motifs with blaOXA-23. We designated this PBP as PbpM and sometime referred as Pbp5 or DacB but has no similarity to *E. coli* PBP5 or DacB. The primers for blaC (sometime referred as blaA) were made and blaCF= 5’-GCT CAC GCA TCT GGA CAA AC-3’ and blaCR= 5’-CCA TCC AAT CGG TGA GCA GT-3’ with Tm=60^0^C and the product size=454bp. The primers of DacB1 was also made and DacB1F= 5’-ACG CTC ACC CTC AAC AAG TC-3’ and DacB1R= 5’-GTT GAA CCC GTA GTC GAG CA-3’ and Tm=60°C and the product size=594bp. The primers for DacB2 were also made and dacB2F= 5’-CCA AGA TGA ACG CCA AAG CC-33’ and DacB2R= 5’-TCG CAG CCT GAT CCC AAT AC-3’ and the Tm=60°C and the product size=390bp. Thus, after complete sequencing of genome of *Mt*, the position of individual PBP could be assigned using any forward for reverse primers and BLAST-2 search.

## Discussion

*Escherichia coli* possesses five high-molecular-weight (HMW) PBPs, with three (PBP1a, PBP1b, and PBP1c) in class-A and two (PBP2 and PBP3) in class-B. This organism has seven low-molecular-weight (class-C) D,D-carboxypeptidases or D,D-endopeptidases (PBP4, PBP5, PBP6, PBP6b, PBP7, PBP4b, and AmpH) [41–44]. Using BLASTP and BLASTN homology searches, we described the correct nomenclature of *Mt* PBPs. Database suggested a wide distribution of mutations among the different β-lactamases [45]. Mutations affect the specificities to β-lactams hydrolysis. As for example, mutation of Serine-238 or Lysine-240 of blaSHV chromosomal enzyme increased specificities to ceftazidime and cefotaxime respectively as comparable to blaCTXM-9 which could hydrolysis cephalosporin drug efficiently. The ESBLs TEM-155 (Q39K, R164S, E240K) and SHV-105 (I8F, R43S, G156D, G238S, E240K) and a non-ESBL, SHV-48 (V119I) mutations affected drug specificities [53]. The sequence similarity between blaTEM-1 and blaNDM-1 was not found but both hydrolysed the penicillin drugs. The blaOXA-1 favours the hydrolysis of oxacillin than ampicillin but blaOXA-23 and blaOXA-51 prefer to hydrolysis carbapenem drugs. Thus, roles of PBPs in penicillin-binding and penicillin-hydrolysis of few hundred penicillin, cephalosporin and carbapenem drugs has a complex mechanism. The original blaCTX-M and blaGES enzymes were ESBLs that hydrolyse both penicillin and cephalosporin well, but not aztreonam and carbapenem drugs and were inhibited by the newer β-lactamase inhibitors such as avibactam, vaborbactam and relebactam [43]. The AmpC β-lactamases are cephalosporinases encoded on the chromosomes of many of the Enterobacteriaceae where they mediate resistance to early cephalosporins like cephalothin, cefazolin, cefoxitin, ampicillin, amoxicillin, and few β- lactamase inhibitor-β-lactam like amoxicillin-cavulinate combinations [54]. It was suggested that cefotaxime, ceftriaxone, ceftazidime, cefepime, cefuroxime, piperacillin, and aztreonam were weak inducers and weak substrates for AmpC class enzymes. We pinpointed here that *M. tuberculosis* penicillin-resistance was mediated by four chromosomal AmpC-like β-lactamases whereas other authors neglected such enzymes in their analysis and accepted only one AmpH enzyme in *Mt*. The analysis corrected that blaAmpH3 is more like blaDHA than blaAmpC (table-3). Kumar et al recently described PonA1/A2 enzymes as best β-lactams binding and hydrolysis although data presented the opposite fact (PonA1/A2= <200 M^-1^S^-1^ vs and AmpH=>4000 M^-1^S^-1^) [8]. We detected conserved SxxK motif in PonA1/A2, PBPA/B/C and also AmpH1, AmpH2, AmpH3 and AmpH4 of *Mt* as well as *E. coli* blaAmpC and *P. aeruginosa* blaDHA β-lactamases (figure-2). Our interpretation was that the major β-lactamase activities in *Mt* was regulated by AmpH β-lactamases which had similarity to blaAmpC, blaDHA, blaACC and blaCMY. But blaAmpH3 has no homology to blaAmpC but has homology to blaDHA enzyme. Still question may arise among the four AmpH proteins characterized which would be more AmpC character? Surely, AmpH1 and AmpH4 will have more class-C β- lactamase character with wider homologies whereas functions of AmpH2 and AmpH3 will be confirmed after laboratory testing with purified enzymes. Problem is if you have a low activity then you will not get a good hydrolysis of penicillin. On the other hand, the high concentrated enzyme preparation (cloned in plasmids and over-expressed or induced expressed) gave false data of β-lactam hydrolysis in a radioactive or fluorescence-tagged antibiotic. Our guess the hydrolysis by many PBPs like PonA1 and PonA2 were happened due to over-expressed enzyme and in vivo effects of those proteins for penicillin drug resistance was not surely happening!

Further, no *Mt* PBP with homology to *E. coli* Pbp3 was found and some author described *Mt* PbpB as Pbp3 in the literature [55, 56]. Similarly, AmpH or DacB1 described as PBP4 in *Mycobacterium* database. But correct PBP4 enzyme was located in the genomes of *Mt* strains OA115DS, J09400698, CCDC5079 and ATCC-35743 whereas in standard strain H37Rv reported as conserved protein (AN:AL123456) [57, 58]. The carbapenemase homology detected in blaOXA-23-like PBPA/B enzymes (protein ids. CCP42738 and CCP44940) and blaOXA-58 related blaMec_N enzyme (protein id. CCP45666) whereas MBLs like blaVIM, blaIMP and blaNDM homologies could not be found (table-2). The ampE, ampD, ampR and ampDE virulence genes specifically in *Pseudomonas aeruginosa* suggested as the regulators of induction and de-repression of AmpC genes whereas DacB was also suggested as an inducer. The ampR located upstream of AmpC gene and was a LysR-type regulator. Interestingly we located few LysR-like locus in *Mt* chromosome, upstream of AmpH1, AmpH2, AmpH3 and AmpH4 genes (data not shown). Plasmid-mediated AmpC gene was also found recently and blaZ of *S. aureus* has 30% homology to blaC β-lactamse of *Mt* and blaCARB gene of *V. parahaemolyticus* (data not shown). As comparison, an induced blaAmpC gave MIC90 250 µg/ml for cefotaxime but it reduced to MIC90 0.06 µg/ml without such induction [59]. Interestingly BLASTN search with *Mt* genome did not match greatly with an inducer (ampR) but three loci were detected with GC-rich minor similarities. But we did not find any ampR protein at the upstream of AmpH1, AmpH2, AmpH3 and Amp4 genes in *Mt*. The *Mt* genome had numerous TFs genes like LuxR, ArcR, GntR, LytR, TetR and FadR and thus activation may possible in trans. Thus, our point of view was that AmpH genes were expressed in low amount as compared to inducible AmpC gene found in *P. aeruginosa* or plasmid-mediated AmpH genes [47].

Bhakta and Basu purified a truncated PonA2 with huge binding to amoxicillin and cefotaxime (K2/K=50000 M^-1^S^-1^) but no comparable data found with *Mt* other PBPs [42]. Nadler JP et al described the some PBP-like similarities in *Mt* but disclosed MT0202 (AAK44422; AE000516.2, nt. 223676-224776), MT0501 (AAK44724; AE000516.2, nt. 573158-574513) and MT1477 (AAK45741; AE000516.2, nt. 1611290-1612105) had no L,D-transpeptidase activities to make 3->3 linkage in *Mt* instead 4->3 linkage of peptidoglycan cross linking but MT0125 (AAK44348; AE000516.2; nt.140258-140890) did well. However, such enzymes had no similarities to β-lactamases [60, 61]. Thus, D,D-carboxypeptidase PBPs are different enzyme and have found good penicillin-binding. The β-lactams hydrolysis by concentrated enzyme (over-expressed and purified) in vitro may not be comparable to its *in vivo* function.

*Streptococcus pneumoniae* DacB acts as an L,D-carboxypeptidase towards the tetrapeptide L- Ala-D-iGln-L-Lys-D-Ala of the peptidoglycan stem, with Km and kcat values of 2.84 mM and 91.49 S^-1^ respectively [62]. Satishkumar et al demonstrated that PBP4 of *Staphylococcus aureus* could not be considered to be a classical mediator of β-lactam resistance and our data analysis supported the similar view in *Mt* [63, 64]. However, *S. aureus* MecA and PBP4 both had crucial role in penicillin-drug resistance. Recently, an inhibitor of PBP4 was discovered to act as antibacterial drug [65]. Surprisingly, there was report that mutations in PBP4 (V223I and A617T) caused decreased susceptibility to penicillin and carbapenem in *Enterococcus faecalis* [57]. A Penicillin-binding-protein-2 (PenA) was shown to catalyse head-to-tail macrolactamization of non-ribosomal peptides whereas a similar enzyme, SurE had higher chains length peptides specificity [66]. Interestingly, in *Mycobacterium* PBP2 had two isomers (PBPA and PBPB) to generate early cephalosporin resistance but in *Neisseria gonorrhoeae* five such isomers were detected [67]. Thus, identification of four blaAmpH isomers regulating in *Mt* was not surprising. The PonA1/A2 conserved SxxN and SxN penicillin-binding motifs were shown at the carboxy-terminus but KxG conserved motif in PonA1 was changed to “PMS” in PonA2. The conserved SxxK, SxN and KxG amino acid sequences in PBPA, PBPB and PBPC (MecA_N) of *M. tuberculosis* also demonstrated as compared with *E. coli* and *S. aureus* PBP2 (see, S5). The conserved SxxK sequence in AmpH1, AmpH2, AmpH3 and AmpH4 *M. tuberculosis* β-lactamases were found but no KxG and SxN motifs were detected. As there were no much conserved domain among four AmpH proteins, we predicted extensive small deletions and mutation during selection of those conserved AmpC and DHA β-lactamase motifs. Thus, even we did not find any similarity between ponA1/A2 with any 24 β-lactamases, still some PBP-motifs indeed was present and widely β-lactam hydrolysis were detected by different workers [8]. Interestingly, Wivagg et al demonstrated that some mutations in the PonA2 gene or inactivation of the gene gave penicillin drug resistance instead penicillin sensitivity demonstrating penicillin-binding and penicillin-cleavage were two different functions [68, 69]. The PBP-motif located however, at the carboxy-terminus of the proteins and likely glycosyltransferase activity located at the amino-terminus of PonA1 and PonA2. Similarly, Kieser et al demonstrated using similar transposon-tagged mutagenesis approach that peptidoglycan synthesizing enzymes like PonA1 plus PonA2 (peptidoglycan transglycosylase and 4,3 transpeptidase) and IdtB (3,3 transpeptidase) proteins have *Mycobacterium* specific mechanism to make peptidoglycan and binding of penicillin varied considerably among isomers [70]. The *Neisseria gonorrhoeae* PonA1 gene mutation was also shown to cause penicillin drug resistance [71]. The *V. prarahaemolyticus* carbenicillin- hydrolysing activity was associated with chromosomal blaCARB gene but PonA1 gene was also located. Dekhil and Mardassi showed recently the important of PonA1, Rv0197, esxK and eccC2 genes mutations in *Mt* sub-lineages L4.3 evolution [72]. Thus, PonA1/A2 have roles other than penicillin-binding but carbapenem hydrolysis is under question.

Interestingly, we detected a relation of ISKpn6 transposase and blaAmpH4 gene as 17nt sequence perfectly matched. We BLASTN searched *Mt* chromosomes with 17nt oligonucleotides and selected many *Mt* chromosomes at the blaAmpH4 locus whereas all AmpH4 proteins appeared conserved and only two mutations detected for *Mt* strain 5521 AmpH4 protein (figure-12). On the other hand, many bacteria had the one AmpH gene and such proteins had high similarities although mutations were detected in different Genus of bacteria (figure-11). Surely, we suggested a role of IS1182-like ISKpn6 transpose in the genome rearrangement of *Mycobacterium* and such deletions, mutations and rearrangements could cause new gene or also could inactivate another gene as happened in *Mt* FDAARGOS_756 AmpH4 gene (data not shown).

MβL enzymes alone occupy class B which is subdivided into B1, B2 and B3 sub-classes evolution events and require bivalent metal ions such as Zn^2+^ and Mg^2+^ for their activity. The catalytic site contained a conserved motif (HxHxDH) and more downstream residues (H196 and H263), appeared to be ancestrally shared by the superfamily of MβL fold proteins. More than 34,000 proteins with diverse biological functions including β-lactamases, nucleases, ribonucleases, lactonases, glyoxalases, hydrolases, Aryl sulfatases, Alkylsulfatases, and others were reported in all organisms. In bacteria, more than 325 variants grouped into 63 MβL types and divided into three sub-groups: subgroup B1 (e.g., NDM-1, VIM-2, and IMP-1); B2 (e.g., CphA1, CphA7, and ImiS) and B3 (e.g., GOB-13, LRA-1, and CAR-1). But such conserved motifs were lacking in PBPs of *Mt* [73]. Truly, MBL mechanism in *Mt* was still elusive. The AmpC is originally detected in chromosome (*E. cloacae, S. marcescens* and *P. aeruginosa*) but now mobilised into variety of plasmids. The AmpR gene regulates AmpC genes negatively. Certain β-lactams induce the production of cell-wall degradation products (eg, *N*- acetylglucosamine-1,6-anhydro-*N*-acetylmuramic acid oligopeptides) which compete with uridine diphosphate (UDP)–*N*-acetylmuramic acid peptides for binding to AmpR. With decreased UDP-*N*-acetylmuramic acid peptide binding to AmpR, results in higher production of AmpC enzymes [74]. Further ampD and ampG regulates such membrane oligopeptides production and transport. The *E. coli* class-C β-lactamases like blaCMY, bla*F*OX, blaDHA, blaACC, blaACT, blaMIR, blaDHA, and blaMOX, produced in plasmids leading to resistance to expanded-spectrum cephalosporins.

The DacD gene was cloned which was also known as PBP6B and was located in *E. coli* (AN: LN832404), *K. pneumoniae* (AN: NZ_CP101563) and *S. enterica* (AN: NZ_CP129206) genomes but such gene was not detected in *M. tuberculosis* (AN: AL123456), *M. leprae* (AN: AL450380), *S. aureus* (AN; BX571856), *S. pneumoniae* (AN: NZ_CP107038) and *C. diphtheriae* (AN: NZ_CP040523) genomes and also little homology detected with both *V. cholerae* chromosomes (AN: NZ_CP028892 for Ch-1 and NZ_CP012998 for Ch-2). Thus, we greatly compared the presence of all PBPs in *M. tuberculosis*. We disclosed four class-C β- lactamases in the hydrolysis of cephalosporin drugs which never described well although presence of AmpH and AmpC class enzyme activity indirectly reported in the NCBI database. The *M. leprae* was a different entity and caused leprosy in India and many other parts of Asia.

*M. leprae* has genetic instability [75–77]. We analysed in details the localization and characterization of *AmpH* genes in *M. leprae* as compared to *M. tuberculosis* and *S. aureus*. All AmpH1, AmpH2 and AmpH3 genes were pseudogenes in *M. leprae* but could be detected by BLAST-N search (figure-8). But we did not able to detect *AmpH4* gene fragment in *M. leprae* suggesting complete deletion. More interestingly, *blaC* gene was also silent in *M. leprae* whereas mmpL5 important mediator of multi-drug resistance was also missing in *M. leprae* [52]. The PonA1, PonA2, PbpA, PbpB and MecA_N proteins were very similar to *M. tuberculosis* (90% similarities) and penicillin resistance was mediated by these enzymes in *M. leprae* [78–80]. Interestingly, similar to TB, Leprosy treatment was never suggested by any β- lactam drug! By virtue of analysis, however, we predicted that *M. leprae* would be more sensitive to penicillin drugs. The analysis of bacterial PBP1B or PonA2 protein multi-alignment and phylogenetic analysis found *M. leprae* also association with *S. aureus* while *M. tuberculosis, M. bovis, M. asiaticum, and M. canittii* were in one group whereas *M. avium, M. colombiens* and *M. intracellulare* in a different group (figure-10). Thus, sequence similarities among the *bla*AmpH genes of different *Mycobacterium* species found well but quite different than *E. coli*-related Gram-negative bacteria or Gram-positive bacteria like *Vibrio*, *Streptococcus* and *Staphylococcus* species. Surely, now over-expression-purification and characterization of AmpH proteins of *M. tuberculosis* to develop a new drug against TB is a priority area of research. Further, comparison of PBPs in other important pathogens was reported [81, 82]. The phage therapy protocols were also progressing in mycobacterial diseases [83, 84]. The phyto-drugs were another remedy protocol for TB and *Mt* RNA polymerase was inhibited by a terpentine-polybromophenol from *Cassia fistula* (golden showers) bark [85, 86]. We discussed the PENEM drug resistance in *Mt* but multi-resistance against other TB specific drugs (rifampicin, dapsone, bedaquiline, isoniazid, moxifloxacin) were more complex [87–90].

## Conclusion

So far ponA1, PonA2, PbpA, PbpB, DacB1 and DacB2 were implicated in penicillin-binding and hydrolysis in *M. tuberculosis.* Using bioinformatics BLAST homology approach, we first time showed that blaAmpH type four class-C β-lactamases similar to blaAmpC, blaDHA, blaACC and blaCMY control the cephalosporin hydrolysis in *Mt* and such genes appeared as pseudogenes in *M. leprae.* The *Mt* AmpH enzymes had better homologies with *V. parahaemolyticus* or *Yerdinia pekkanenii* AmpH enzyme than *E. coli* AmpH enzyme. The blaC class-A β-lactamase has similarity to *S. aureus* blaZ β-lactamase as well as blaCARB gene of *V. parahaemolyticus* but such gene also inactivated in *M. leprae* including an important drug efflux gene called mmpL5. Thus, we showed a different new mechanism of PENEM drug resistance in *M. tuberculosis* and *M. leprae.* All primers for *Mt* PBPs were made to check the transcription of individual genes by RT-PCR as well as to check chromosomal location by BLASTN search after WGS to avoid the concurrent mistakes in the nomenclature of *Mt* PBPs. Interestingly, we showed that ISKpn6 transposase gene was involved in the genesis of *bla*AmpH4 gene suggesting the mechanism of chromosomal genesis of β-lactamases likely under high dose of penicillin and cephalosporin drugs.

## Acknowledgement

We thank NCBI for free database and CLUSTAL-Omega free software. AKC is retired professor of Biochemistry. AKC thanks Prof. Bidyut Bandhopadhyay, Principal of OIST for his interest in Mycobacterium research. AKC also thanks Prof. Sujoy Dasgupta, retired chairman of Microbiology department, Bost Institute, Kolkata for suggestion on tuberculosis research.

## References

[1] Chakraborty AK. Current status and unusual mechanism of multi-resistance in Mycobacterium tuberculosis. J Health Med Informatics. 2019; 10(1): 328. Doi: 10.4172/2157-7420.1000328.

[2] Chakaya J, Khan M, Ntoumi F, Aklillu E, Fatima R, Mwaba P, et al. Global tuberculosis report 2020 - reflections on the global TB burden, treatment and prevention efforts. Int J Infect Dis. 2021; 113 Suppl 1, S7– S12. doi: 10.1016/j.ijid.2021.02.107.

[3] World Health Organization. Molecular line probe assays for rapid screening of patients at risk of multidrug-resistant tuberculosis (MDR-TB). Geneva: World Health Organization; 2008.

[4] World Health Organization/IUATLD Global Project on Anti-Tuberculosis Drug Resistance Surveillance. Anti-tuberculosis drug resistance in the world. Publication no. WHO/TB/97.229. Geneva, Switzerland: World Health Organization; 1998.

[5] Palomino JC, Martin A. Drug resistance mechanisms in Mycobacterium tuberculosis. Antibiotics 2014, 3, 317–340.

[6] Soltys MA. The effect of penicillin on Mycobacteria in vitro and in vivo. Tubercle. 1952;33(4):120–125. Doi: 10.1016/S0041-3879(52)80054-6.

[7] Kasik JE, Weber M, Winberg E, Barclay WR. The synergistic effect of dicloxacillin and penicillin G on murine tuberculosis. Am Rev Respir Dis. 1966;94(2):260–261.

[8] Kumar G, Galanis C, Batchelder HR, Townsend CA, Lamichhane G. Penicillin Binding Proteins and b- Lactamases of Mycobacterium tuberculosis: Re-examination of the Historical Paradigm. mSphere, 2022; 7 (1): e00039–22.

[9] Wivagg CN, Bhattacharyya RP, Hung DT. Mechanisms of b-lactam killing and resistance in the context of Mycobacterium tuberculosis. J Antibiot. 2014; 67:645–654. 10.1038/ja.2014.94.

[10] Eric-Sauvage E, Kerff F, Terrak T et al. The penicillin-binding proteins: structure and role in peptidoglycan biosynthesis. FEMS Microbiology Reviews, 2008; 32(2): 234–258. 10.1111/j.1574-6976.2008.00105.x.

[11] Campbell PJ, Morlock GP, Sikes RD et al. Molecular detection of mutations associated with first- and second-line drug resistance compared with conventional drug susceptibility testing of *M. tuberculosis*. Antimicrob Agents Chemother. 2011;55(5), 2032–2041. doi: 10.1128/AAC.01550-10.

[12] Campbell EA, Korzheva N, Mustaev A, Murakami K, Nair S, Goldfarb A and Darst SA. Structural mechanism for rifampicin inhibition of bacterial RNA polymerase. Cell, 2001; 104(6), 901–912.PMID:11290327.

[13] García SR, Otal I, Martín C, Gómez-Lus R, Aínsa JA, et al. Novel streptomycin resistance gene from Mycobacterium fortuitum. Antimicrob Agents Chemother. 2006; 50: 3920–3922.

[14] Sreevatsan S, Pan X, Stockbauer KE et al. Characterization of *rpsL* and *rrs* mutations in streptomycin-resistant *Mycobacterium tuberculosis* isolates from diverse geographic localities. Antimicrob Agents Chemother. 1996; 40(4), 1024–1026.

[15] Hegde SS, Vetting MW, Roderick SL et al. A fluoroquinolone resistance protein from *Mycobacterium tuberculosis* that mimics DNA. Science, 2005; 308(5727), 1480–1483. PMID:15933203.

16. WHO. Multidrug and extensively drug-resistant TB (M/XDR-TB): 2010 global report on surveillance and response, World Health Organization, Geneva, Switzerland, http://whqlibdoc.who.int/ publications/2010/9789241599191_eng.pdf.

[17] Islam MM, Tan Y, Hameed HMA, Liu Z, Chhotaray C, et al. Detection of novel mutations associated with independent resistance and cross-resistance to isoniazid and prothionamide in Mycobacterium tuberculosis clinical isolates. Clin Microbiol Infect. 2018; Dec 22. pii: S1198-743X(18)30794-8. doi: 10.1016/j.cmi.2018.12.008.

[18] Almeida Da Silva, P.E.; Palomino, J.C. Molecular basis and mechanisms of drug resistance in M. tuberculosis: Classical and new drugs. J. Antimicrob. Chemother. 2011; 66: 1417–1430.

[19] Madsen CT, Jakobsen L, Buriankova K et al. (2005) Methyltransferase Erm(37) slips on rRNA to confer atypical resistance in *Mycobacterium tuberculosis*. J Biol Chem. 280(47), 38942–38947. DOI:10.1074/jbc.M505727200.

[20] Zhang, B. et al. Crystal structures of membrane transporter MmpL3, an anti-TB drug target. Cell, 2019;176: 636.e3–648.e3.

[21] Zheng H, Lu L, Wang B, Pu S, Zhang X, Zhu G, Shi W, et al. Genetic basis of virulence attenuation revealed by comparative genomic analysis of Mycobacterium tuberculosis strain H37Ra versus H37Rv. PLoS ONE, 2008; 3 (6): E2375.

[22] Alderwick LJ, Harrison J, Lloyd GS and Birch HL. The Mycobacterial cell wall-peptidoglycan and arabinoglycan. Cold Spring Harbor Perspect Med. 2015; 5: a021113. doi:10.1101/cshperspect.a021113.

[23] He XC, Zhang XX, Zhao JN, et al. Epidemiological trends of drug-resistant tuberculosis in China from 2007 to 2014: A retrospective study. Medicine (Baltimore), 2016;95(15), e3336. PMID:27082586.

[24] Duan Q, Chen Z, Chen C, et al. The Prevalence of drug-resistant tuberculosis in mainland China: An updated systematic review and meta-analysis. PLoS One, 2016; 11(2): e0148041. Doi:10.1371/journal.pone.0148041.

[25] Vilchèze C, Jacobs WR Jr. The isoniazid paradigm of killing, resistance, and persistence in Mycobacterium tuberculosis. J Mol Biol. 2019; pii: S0022-2836(19)30093-2. doi: 10.1016/j.jmb.2019.02.016.

[26] Scorpio A and Zhang Y. Mutations in *pncA* gene encoding pyrazinamidase/nicotinamidase, cause resistance to the antituberculous drug pyrazinamide in tubercle bacillus. Nature medicine, 1996; 2(6): 662–667.

[27] Smith T, Wolff KA and Nguyen L. Molecular Biology of Drug Resistance in *Mycobacterium tuberculosis*. Curr Top Microbiol Immunol. 2013; 374: 53–80. Doi: 10.1007/82_2012_279.

[28] Uchiya K, Takahashi H, Nakagawa T, Yagi T, Moriyama M, Inagaki T, Ichikawa K, Nikai T, Ogawa K. Characterization of a novel plasmid, pMAH135, from Mycobacterium avium subsp. hominissuis. PLoS One. 2015;10(2): e0117797. Doi: 10.1371/journal.pone.0117797.

[29] Matsumoto CK, Bispo PJ, Santin K, Nogueira CL, Leão SC. Demonstration of plasmid-mediated drug resistance in Mycobacterium abscessus. J Clin Microbiol. 2014; 52(5):1727–1729. Doi: 10.1128/JCM.00032-14.

[30] Chakraborty AK. Poor correlation of diversified MDR genes in *Gonococci* plasmids: Does alteration in chromosomal DEGs, PBP2 and target mutations sufficient to widespread multi-resistance in *Neisseria gonorrhoeae*? J Health Med Informat. 2018; 9: 310. Doi: 10.4172/2157-7420.1000310.

[31] Chakraborty AK, Maity M, Patra S, Mukherjee S and Mandal T. Complexity, heterogeneity and mutational analysis of antibiotic inactivating acetyl transferases in MDR conjugative plasmids conferring multi-resistance. Res Rev: J Microbiol Biotechnol. 2017; 6(2): 28–43.

[32] Das S, Datta S, Dey M, Poria K, Chakraborty AK. Green Chemistry: Phyto-antibiotics, A green antibiotics, isolated from medicinal plant of West Bengal targeting Multi-drug Resistant bacteria”. Acta Scientific Medical Sciences, 2022; 6(5): 127–143.

[33] Chakraborty AK. Complexity, heterogeneity, 3-D structures and transcriptional activation of multi-drug resistant clinically relevant bacterial beta-lactamases. Trends Biotechnol-open access.2016; 2(1):1–001.

[34] Jacoby GA. AmpC β-lactamases. Clini Microbiol Rev. 2009; 22(1): 161-182. Doi: 10.1128/CMR.00036-08.

[35] Chakraborty AK, Pradhan S, Das S, Maity M, Sahoo S, Poria K. Complexity of OXA Beta-Lactamases involved in Multi-Resistance. British J Bio-Medical Res. 2019; 3(1): 772–798. Doi: 10.24942/bjbmr.2019.424.

[36] Chakraborty AK, Poira K, Saha D, Halder C, Das S. (2018) Multidrug- Resistant Bacteria with activated and diversified MDR Genes in Kolkata Water: Ganga Action Plan and Heterogeneous Phyto-Antibiotics tackling superbug spread in India. American J Drug Deli Therapeutics, 2018; 5(1): 2. Doi: 10.13140/RG.2.2.17947.52001.

[37] Fleming A. On the antibacterial action of cultures of a penicillium, with special reference to their use in the isolation of B. influenzae 1929. Bull World Health Organ. 2001;79:780–790.

[38] Tan SY, Tatsumura Y. Alexander Fleming (1881-1955): Discoverer of penicillin. Singapore Med J. 2015;56(7):366–7. doi: 10.11622/smedj.2015105.

39. Lynch JP 3rd, Clark NM, Zhanel GG. Evolution of antimicrobial resistance among Enterobacteriaceae (focus on extended spectrum beta-lactamases and carbapenemases). Exp. Opini. Pharmacother, 2013, 14, 199–210.

[40] Ghuysen JM. Serine beta-lactamases and penicillin-binding proteins. Annu Rev Microbiol. 1991; 45:37–67.

[41] Goffin C, Ghuysen JM. Multimodular penicillin-binding proteins: an enigmatic family of orthologs and paralogs. Microbiol Mol Biol Rev. 1998;62(4):1079–1093. doi: 10.1128/MMBR.62.4.1079-1093.1998.

[42] Bhakta S and Basu J. Overexpression, purification and biochemical characterization of a class A high-molecular-mass penicillin-binding protein (PBP), PBP1* and its soluble derivative from Mycobacterium tuberculosis. Biochem J. 2002;361:635–639.

[43] Voladri RK, Lakey DL, Hennigan SH, Menzies BE, Edwards KM, Kernodle DS. Recombinant expression and characterization of the major beta-lactamase of Mycobacterium tuberculosis. Antimicrob Agents Chemother. 1998; 42:1375–1381. 10.1128/AAC.42.6.1375.

[44] Sung MT, Lai YT, Huang CY, Chou LY, et al. Crystal structure of the membrane-bound bifunctional transglycosylase PBP1b from Escherichia coli. Proc Natl Acad Sci USA, 2009; 106: 8824–8829.

[45] Schiffer G, Holtje JV. Cloning and characterization of PBP 1C, a third member of the multi-modular class A penicillin-binding proteins of *Escherichia coli*. J Biol Chem. 1999;274:32031–32039.

[46] Diene SM, Pontarotti P, Azza S, Armstrong N, et al. Origin, Diversity, and Multiple Roles of Enzymes with Metallo-β-Lactamase Fold from Different Organisms. Cells. 2023;12(13):1752. doi: 10.3390/cells12131752.

[47] González-Leiza SM, de Pedro MA, Ayala JA. AmpH, a bifunctional DD-endopeptidase and DD-carboxypeptidase of Escherichia coli. J Bacteriol. 2011;193(24):6887–6894. doi: 10.1128/JB.05764-11.

[48] Henderson TA, Young KD, Denome SA, Elf PK. AmpC and AmpH, proteins related to the class C beta-lactamases, bind penicillin and contribute to the normal morphology of Escherichia coli. J Bacteriol. 1997;179(19):6112–6121. doi: 10.1128/jb.179.19.6112-6121.1997.

[49] Corpet F. Multiple sequence alignment with hierarchical clustering. Nucl. Acids Res. 1988; 16(22): 10881–10890.

[50] Sievers F, Wilm A, Dineen DG, Gibson TJ, Karplus K, et al. Fast, scalable generation of high-quality protein multiple sequence alignments using Clustal Omega. Molecular Systems Biology. 2011; 7, Article number: 539. doi:10.1038/msb.2011.75.

[51] Altschul SF, Gish W, Miller W, Myers EM, Lipman DJ. Basic local alignment search tool. J Mol Biol. 1990; 215:403–410. DOI: 10.1016/S0022-2836(05)80360-2.

[52] Cole ST, Eiglmeier K, Parkhill J et al. Massive gene decay in the leprosy bacillus. Nature, 2001; 409: 1007–1011

[53] Tamma PD, Aitken SL, Bonomo RA, Mathers AJ, van Duin D, Clancy CJ. Infectious Diseases Society of America Guidance on the Treatment of AmpC β-Lactamase-Producing Enterobacterales, Carbapenem-Resistant Acinetobacter baumannii, and Stenotrophomonas maltophilia Infections. Clin Infect Dis. 2022;74(12):2089–2114. doi: 10.1093/cid/ciab1013.

[54] Jaurin B and Grunström T. *ampC* cephalosporinase of *Escherichia coli* K-12 has a different evolutionary origin from that of β-lactamases of the penicillinase type. Proc Natl Acad Sci USA. 1981;78:4897–4901.

[55] Lim D and Strynadka NC. Structural basis for the beta-lactam resistance of PBP2a from methicillin-resistant *Staphylococcus aureus*. Nat Struct Biol. 2002; 9:870–876.

[56] Pares S, Mouz N, Petillot Y, Hakenbeck R et al. X-ray structure of *Streptococcus pneumoniae* PBP2x, a primary penicillin target enzyme. Nat Struct Biol. 1996;3:284–289.

[57] Rice LB, Desbonnet C, Tait-Kamradt A, Garcia-Solache M, Lonks J, Moon TM, D’Andréa ÉD, Page R, Peti W. Structural and Regulatory Changes in PBP4 Trigger Decreased β-Lactam Susceptibility in Enterococcus faecalis. mBio. 2018;9(2):e00361–18. doi: 10.1128/mBio.00361-18.

[58] Basuino L, Jousselin A, Alexander JAN, Strynadka NCJ, Pinho MG, Chambers HF, Chatterjee SS. PBP4 activity and its overexpression are necessary for PBP4-mediated high-level β-lactam resistance. J Antimicrob Chemother. 2018;73(5):1177–1180. doi: 10.1093/jac/dkx531.

[59] Poirel L, Guibert M, Girlich D, et al. Cloning, Sequence Analyses, Expression, and Distribution of *ampC ampR* from *Morganella morganii* Clinical Isolates. Antimicrob Agents chemther. 1999; 43:769–776. Doi: 10.1128/aac.43.4.769.

[60] Nadler JP, Berger J, Nord JA, Cofsky R, Saxena M. Amoxicillin-clavulanic acid for treating drug-resistant *Mycobacterium tuberculosis*. Chest. 1991;99:1025–1026.

[61] Gupta R, Lavollay M, Mainardi JL, Arthur M, Bishai WR, Lamichhane G. The *Mycobacterium tuberculosis* protein LdtMt2 is a nonclassical transpeptidase required for virulence and resistance to amoxicillin. Nature Medecine. 2010;16:466–469.

[62] Zhang J, Yang YH, Jiang YL, Zhou CZ, Chen Y. Structural and biochemical analyses of the Streptococcus pneumonia L,D-carboxypeptidase DacB. Acta Crystallogr D Biol Crystallogr. 2015;71(Pt 2):283–92. doi: 10.1107/S1399004714025371.

[63] Satishkumar N, Alexander JAN, Poon R, Buggeln E, Argudín MA, Strynadka NCJ, Chatterjee SS. PBP4- mediated β-lactam resistance among clinical strains of Staphylococcus aureus. J Antimicrob Chemother. 2021;76(9):2268–2272. doi: 10.1093/jac/dkab201.

[64] da Costa TM, de Oliveira CR, Chambers HF, Chatterjee SS. PBP4: A New Perspective on *Staphylococcus aureus* β-Lactam Resistance. Microorganisms. 2018;6(3):57. doi: 10.3390/microorganisms6030057.

[65] Young M, Walsh DJ, Masters E, et al. Identification of *Staphylococcus aureus* Penicillin Binding Protein 4 (PBP4) Inhibitors. Antibiotics (Basel). 2022;11(10):1351. doi: 10.3390/antibiotics11101351.

[66] Matsuda K, Fujita K, Wakimoto T. PenA, a penicillin-binding protein-type thioesterase specialized for small peptide cyclization. J Ind Microbiol Biotechnol. 2021;48(3-4):kuab023. doi: 10.1093/jimb/kuab023.

[67] Ohnishi M, Watanabe Y, Ono E, Takahashi C, Oya H, Kuroki T, Shimuta K, Okazaki N, Nakayama S, Watanabe H. Spread of a chromosomal cefixime-resistant penA gene among different Neisseria gonorrhoeae lineages. Antimicrob Agents Chemother. 2010;54(3):1060–1067. doi: 10.1128/AAC.01010-09.

[68] Wivagg CN, Wellington S, Gomez JE, Hung DT. Loss of a Class A Penicillin-Binding Protein Alters β- Lactam Susceptibilities in Mycobacterium tuberculosis. ACS Infect Dis. 2016;2(2):104–110. doi: 10.1021/acsinfecdis.5b00119.

[69] Deshpande D, Srivastava S, Chapagain M, et al. Ceftazidime-avibactam has potent sterilizing activity against highly drug-resistant tuberculosis. Sci Adv. 2017;3(8):e1701102. doi: 10.1126/sciadv.1701102.

[70] Hett EC, Chao MC, Rubin EJ. Interaction and modulation of two antagonistic cell wall enzymes of mycobacteria. PLoS Pathog. 2010;6(7):e1001020. doi: 10.1371/journal.ppat.1001020.

[71] Shigemura K, Shirakawa T, Massi N, et al. Presence of a mutation in ponA1 of Neisseria gonorrhoeae in numerous clinical samples resistant to various beta-lactams and other, structurally unrelated, antimicrobials. J Infect Chemother. 2005;11(5):226–30. doi: 10.1007/s10156-005-0403-1.

[72] Dekhil N and Mardassi H. Genomic changes underpinning the emergence of a successful *Mycobacterium tuberculosis* Latin American and Mediterranean clonal complex. Front Microbiol. 2023;14: 1159994. doi: 10.3389/fmicb.2023.1159994.

[73] Bebrone C. Metallo-β-lactamases (classification, activity, genetic organization, structure, zinc coordination) and their superfamily. Biochem Pharmacol. 74: 1686–1701. Doi: 10.1016/j.bcp.2007.05.021

[74] Kocaoglu O and Carlson EE. Profiling of β-lactam selectivity for penicillin-binding proteins in Escherichia coli strain DC2. Antimicrob Agents Chemother. 2015;59(5):2785–90. doi: 10.1128/AAC.04552-14.

[75] Kwon HH, Tomioka H, Saito H. Distribution and characterization of beta-lactamases of mycobacteria and related organisms. Tuber Lung Dis. 1995;76(2):141–8. doi: 10.1016/0962-8479(95)90557-x.

[76] Kasik JE and Peacham L. Properties of beta-lactamases produced by three species of mycobacteria. Biochemical J. !968;107: 675–682. doi: 10.1042/bj1070675.

[77] Sugawara-Mikami M, Tanigawa K, Kawashima A et al. Pathogenicity and virulence of Micobacterium leprae. Virulence, 2022; 13:1985–2011. Doi: 10.1080/21505594.2022.2141987.

[78] Jones CH, Ruzin A, Tuckman M et al. Protocol for Rapid Determination of TEM- and SHV-Type Extended-Spectrum β-Lactamases in Clinical Isolates and Identification of the Novel β-Lactamase Genes *bla*_SHV-48_, *bla*_SHV-105_, and *bla*_TEM-155_. Antimicrob Agents Chemother. 2009; 53:977–9986.

[79] Halim MZ, Jaafar MM, Teh LK et al. Genome sequencing and annotation of multidrug resistant Mycobacterium tuberculosis (MDR-TB) PR10 strain. Genom Data, 2016; 7: 245–246. DOI:10.1016/j.gdata.2016.01.002.

[80] Blattner FR, Plunkett G 3rd, Bloch CA, Perna NT, Burland V, et al. The complete genome sequence of Escherichia coli K-12. Science, 1997; 277 (5331:1453–1462.

[81] Ropp PA, Hu M, Olesky M, et al. Mutations in *ponA*, the gene encoding penicillin-binding protein 1, and a novel locus, *penC*, are required for high-level chromosomally mediated penicillin resistance in *Neisseria gonorrhoeae*. Antimicrob Agents Chemother. 2002;46:769–777.

[82] Leski TA, Tomasz A. Role of penicillin-binding protein 2 (PBP2) in the antibiotic susceptibility and cell wall cross-linking of *Staphylococcus aureus*: evidence for the cooperative functioning of PBP2, PBP4, and PBP2A. J Bacteriol. 2005;187: 1815–1824.

[83] Guerrero-Bustamante CA, Dedrick RM, Garlena RA, Russell DA, Hatfull GF. Toward a phage cocktail for tuberculosis: susceptibility and tuberculocidal action of mycobacteriophages against diverse Mycobacterium tuberculosis strains. mBio, 2021; 12: e00973–21. Doi: 10.1128/mBio.00973-21.

[84] Froman S, Will DW, Bogen E. Bacteriophage active against virulent Mycobacterium tuberculosis. I. Isolation and activity. Am J Public Health Nations Health, 1954;44:1326–1333. 10.2105/AJPH.44.10.1326.

[85] Jiang RH, Xu HB, Fu J. Outcomes of Chinese herb medicine for the treatment of multidrug-resistant tuberculosis: a systematic review and meta-analysis. Complement. Ther Med. 2015; 23(4), 544–554. DOI:10.1016/j.ctim.2015.06.006.

[86] Chakraborty AK, Saha S, Poria K, et al. A saponin-polybromophenol antibiotic (CU1) from Cassia fistula bark targeting RNA polymerase, Current Research Pharmacol & Drug Discovery, 2022; 3:100090. DOI: 10.1016/j.crphar.2022.100090.

[87] Kadura S, King N, Nakhoul M, et al. Systematic review of mutations associated with resistance to the new and repurposed Mycobacterium tuberculosis drugs bedaquiline, clofazimine, linezolid, delamanid and pretomanid. J Antimicrob Chemother. 2020;75(8):2031–2043. doi: 10.1093/jac/dkaa136.

[88] Xu Y, Jia H, Huang H, Sun Z, Zhang Z. Mutations Found in embCAB, embR, and ubiA Genes of Ethambutol- Sensitive and -Resistant Mycobacterium tuberculosis Clinical Isolates from China. Biomed Res Int. 2015;2015: 951706. doi: 10.1155/2015/951706.

[89] Zheng R, Li Z, He F, Liu H, Chen J, Chen J, Xie X, Zhou J, Chen H, Wu X, et al. Genome-Wide Association Study Identifies Two Risk Loci for Tuberculosis in Han Chinese. Nat. Commun. 2018; 9:4072.

[90] Matsumoto CK, Bispo PJ, Santin K, Nogueira CL, Leão SC. Demonstration of plasmid-mediated drug resistance in Mycobacterium abscessus. J Clin Microbiol. 2014; 52(5):1727–9. doi: 10.1128/JCM.00032-14.

